# ASO-mediated knockdown of GPNMB in mutant-*GRN* and *Grn*-deficient peripheral myeloid cells disrupts lysosomal function and immune responses

**DOI:** 10.1101/2024.07.22.604676

**Authors:** Rebecca L. Wallings, Drew A. Gillett, Hannah A. Staley, Savanna Mahn, Julian Mark, Noelle Neighbarger, Holly Kordasiewicz, Warren D. Hirst, Malú Gámez Tansey

**Author notes:** DaCapo Brainscience, 700 Main Street, Cambridge, 02139, MA, USA.

## Abstract

**Background:** Increases in GPNMB are detectable in FTD-*GRN* cerebrospinal fluid (CSF) and post-mortem brain, and brains of aged *Grn*-deficient mice. Although no upregulation of GPNMB is observed in the brains of young *Grn*-deficient mice, peripheral immune cells of these mice do exhibit this increase in GPNMB. Importantly, the functional significance of GPNMB upregulation in progranulin-deficient states is currently unknown. Given that GPNMB has been discussed as a potential therapeutic target in *GRN*-mediated neurodegeneration, it is vital for the field to determine what the normal function of GPNMB is in the immune system, and whether targeting GPNMB will elicit beneficial or deleterious effects.

**Methods:** The effects of GPNMB knock-down via antisense oligonucleotide (ASO) were assessed in peripheral blood mononuclear cells (PBMCs) from 25 neurologically healthy controls (NHCs) and age- and sex-matched FTD-*GRN* patients, as well as peritoneal macrophages (pMacs) from progranulin-deficient (*Grn*^-/-^) and B6 mice. Lysosomal function, antigen presentation and MHC-II processing and recycling were assessed, as well as cytokine release and transcription.

**Results:** We demonstrate here that ASO-mediated knockdown of GPNMB increases lysosomal burden and cytokine secretion in FTD-GRN carrier and neurologically healthy controls (NHCs) monocytes. ASO-mediated knockdown of GPNMB in *Grn*-deficient macrophages decreased lysosomal pan-cathepsin activity and protein degradation. In addition, ASO-mediated knockdown of GPNMB increased MHC-II surface expression, which was driven by decreased MHC-II uptake and recycling, in macrophages from *Grn*-deficient females. Finally, ASO-mediated knockdown of GPNMB dysregulated IFNγ-stimulated cytokine transcription and secretion by mouse macrophages due to the absence of regulatory actions of the GPNMB extracellular fragment (ECF).

**Conclusions:** Our data herein reveals that GPNMB has a regulatory effect on multiple immune effector functions, including capping inflammation and immune responses in myeloid cells via secretion of its ECF. Therefore, in progranulin-deficient states, the drastic upregulation in GPNMB transcript and protein may represent a compensatory mechanism to preserve lysosomal function in myeloid cells. These novel findings indicate that targeted depletion in FTD-*GRN* would not be a rational therapeutic strategy because it is likely to dysregulate important immune cell effector functions.

## Background

Progranulin (PGRN) is a secreted glycoprotein encoded by the *GRN* gene that is internalized via sortilin and prosaposin receptor-binding and taken to the lysosome where it is cleaved by intracellular cathepsins into its seven subunits called granulins (*1–3*). Homozygous *GRN* mutations cause the lysosomal storage disorder (LSD) neuronal ceroid lipofuscinosis (*4*), whilst heterozygous mutations are associated with frontotemporal dementia (FTD-*GRN*) (*5–7*). FTD-*GRN* is responsible for 5-20% of familial FTD cases, and 1-5% of sporadic FTD cases, with around 70 pathogenic mutations identified thus far (*8*). As well as FTD and LSDs, specific *GRN* variants have also been implicated in both Alzheimer’s Disease (AD) (*9–11*) and Parkinson’s Disease (PD) (*12*). Importantly, the effects of these pathogenic variants have been reported to dysregulate PGRN expression through nonsense mediated decay, with a decrease in *GRN* mRNA and PGRN protein in patient samples across these diseases, including brain, peripheral blood mononuclear cells (PBMCs) and serum (*13–16*). A loss of PGRN expression, and therefore function, is associated with increased risk for neurodegenerative diseases, and understanding mechanisms associated with GRN deficiency may aid in future therapeutic development.

In the brain, PGRN is generally viewed as neuroprotective; although the precise mechanisms are unclear, PGRN has been shown to modulate neuroinflammation and reduce brain volume loss in stroke and head injury models (*17, 18*), and was also reported to prevent dopaminergic neuron loss in an MPTP model of PD (*19*). Lysosomal dysregulation is exacerbated with age in the *Grn* -/- mouse brain, leading to neuroinflammation, synaptic loss, and decreased markers of oligodendrocytes, myelin, and neurons (*20*). However, our group has more recently reported that the effects of *GRN* mutations and deficiencies extend beyond the brain, with loss of PGRN leading to early disruption in the crosstalk between the central nervous system (CNS) and the peripheral immune system which may contribute to neurodegeneration (*21*). In the serum from FTD-*GRN* carriers increased levels of soluble Cluster of Differentiation 163 (CD163) and CC Chemokine Ligand 18 (CCL18) have been detected relative to healthy controls, suggesting increased peripheral macrophage activity (*22*). In addition, lipopolysaccharide binding protein (LBP) levels correlated with white-matter changes in the frontal lobe and Clinical Dementia Rating-FTLD sum of boxes scores of FTD*-GRN* patients, indicating that peripheral immune activity may reflect region-specific brain changes and clinically-relevant behavior. Furthermore, when macrophages isolated from *Grn* -/- mice were challenged with lipopolysaccharide (LPS), they displayed altered levels of *Mcp-1, Il-12p40*, *Tnf,* and *Il-10* transcripts relative to B6 controls (*23*). Glycoprotein non-metastatic B (GPNMB), a transmembrane type I protein that is highly expressed in both macrophages and microglia, and is thought to regulate the innate immune response (*24*) and is one of the most upregulated transcripts and proteins in PGRN-deficient states (*20*). Specifically, recent work has identified an increase in GPNMB in *GRN*-FTD patient brain lysate relative to both non-FTD controls and other FTD-related mutations (*20*). In addition, CSF from FTD-*GRN* patients showed increased amounts of GPNMB relative to non-FTD controls (*20*). Furthermore, an age-dependent increase in the expression of GPNMB protein has been reported in *Grn -/-* mice, with brain lysates from 18-month old, but not 3-month old, *Grn -/-* mice exhibiting increased GPNMB expression relative to B6 controls (*20*). This suggests that, in the CNS, GPNMB upregulation may be a component of later stages of disease progression. Importantly, our group demonstrated that GPNMB is dysregulated in peripheral macrophages in the absence of progranulin at a significantly earlier time point than previously reported in the CNS (*25*). Specifically, peritoneal macrophages from 5-to-6-month-old *Grn* -/- mice exhibited significantly increased GPNMB at both the protein and transcript level, highlighting that PGRN deficiency leads to early immune dysregulation in the periphery that pre-empts dysregulation in the CNS.

To understand the mechanisms underlying PGRN-deficiency and neurodegeneration, we have turned our attention to the relationship between PGRN and GPNMB. While it is currently unclear why GPNMB is elevated in human FTD-*GRN* patients and *Grn* -/- mice, GPNMB has been discussed as a potential therapeutic target in PGRN-mediated neurodegeneration (*21*). Therefore, it is vital for the field to firmly establish whether targeting GPNMB will elicit beneficial or deleterious effects. One intriguing possibility is that upregulation of GPNMB expression is driven by lysosomal dysfunction. This idea is supported by previous reports that lysosomal stress causes upregulation of GPNMB in macrophages (*26*). GPNMB is also elevated in the *substantia nigra* of patients with PD (*27*), a neurodegenerative disease increasingly linked to lysosome dysfunction (*28*). Moreover, chemical (*27*) or genetic inhibition (*29*) of β-glucocerebrosidase activity, leading to perturbed lysosomal function, also causes increased expression of GPNMB. Importantly, PGRN deficiency reduces glucocerebrosidase activity (*30, 31*), suggesting that the lysosomal dysfunction could be a proximal cause of GPNMB upregulation in *Grn* -/- mice and FTD-*GRN* patients.

It has also been suggested that GPNMB upregulation may occur in myeloid cells as a means to modulate or cap inflammation (*24*). GPNMB has been reported to have an important anti-inflammatory role and is vital for disease resolution. For example, RAW264.7 cells overexpressing GPNMB showed diminished levels of IL-6 and IL-12 cytokine release after IFN-γ/LPS treatment when compared to controls (*32*), and primary human periodontal ligament cells (hPDLCs) overexpressing GPNMB showed decreased TNF and IL-12 release after LPS treatment relative to controls (*33*). Given the link between inflammation and PGRN deficiency as previously discussed, it is possible, therefore, that GPNMB may be upregulated to help dampen inflammation.

The aim of this study is to assess the effects of GPNMB knock-down via antisense oligonucleotides (ASOs) on both lysosomal function and immune cell responses in peripheral macrophages from 5-to-6-month-old *Grn* -/- mice and PBMCs from FTD-*GRN* patients in order to ascertain why GPNMB is markedly upregulated in the absence of PGRN in peripheral immune cells. Importantly, we will specifically focus on peripheral myeloid cells given our recent findings demonstrating that upregulation of GPNMB in *Grn* -/- mice is detectable much earlier in the peripheral immune system relative to the CNS (*25*). This study will utilize an ASO-based method of knocking down GPNMB, as opposed to genetic knock-out, because genetic ablation of *GPNMB* does not alter synuclein-related pathology (*34*), which we suspect may be due to an upregulation of compensatory genes. pMacs were utilized for the purpose of this study over other primary macrophages routinely used, such as bone-marrow-derived macrophages (BMDMs), the reasons being two-fold; firstly, as pMacs differentiate *in vivo* with no further differentiation required *ex vivo*, as opposed to BMDMs which require *ex vivo* differentiation, it has been argued that pMacs provide a readout of the responsiveness of the innate immune system in the background of the animal from which they are isolated, particularly informative for knockout animals, and is even more imperative when assessing effects of genotype at different ages (*35*). Given that age is an important factor when studying the role of GPNMB increases in PGRN-deficient models (*25*), this argument is of particular importance for this study. Secondly, pMacs are not a homogenous population of cells, but rather a mix of small and large pMacs (SPMs and LPMs, respectively; Sup. Figure 3A). LPMs are resident to the peritoneal cavity and are traditionally thought of as anti-inflammatory, phagocytic and responsible for the presentation of exogenous antigens (*36*). SPMs, on the other hand, are generated from bone-marrow-derived myeloid precursors which migrate to the peritoneal cavity in response to infection, inflammatory stimuli, or thioglycolate, and present a pro-inflammatory functional profile (*36*). We have previously reported differential effects of genetic mutations associated with neurodegeneration on SPM and LPM function (*37, 38*), therefore having an *ex vivo* model in which the effects of genetic mutations or perturbations on different subtypes of myeloid cell is advantageous.

We observed an increase in GPNMB in both FTD-*GRN* patient monocytes and macrophages from 5-to-6-month-old *Grn* -/- mice. When quantifying lysosomal function, no effect of patient group was observed when comparing control-ASO treated PBMCs. However, upon ASO-mediated knock-down of GPNMB, lysosomal function was significantly affected, revealing a significant reduction in healthy lysosomes in FTD-*GRN* patient cells relative to NHCs. Furthermore, upon GPNMB knock-down, a concomitant increase in HLA-DR and MHC-II expression was observed in both patient and *Grn* -/- cells, respectively. This increase in expression was determined to be caused by altered MHC-II uptake and recycling, driven by the lysosomal deficits produced upon GPNMB knock-down. Furthermore, a decrease in stimulation-dependent pro-inflammatory cytokine release was observed in control ASO-treated cells from both FTD-*GRN* patients and *Grn* -/- macrophages from mice, which was ameliorated upon knock-down of GPNMB. Collectively, we conclude that the earliest reported increases in GPNMB in *GRN* mutant/deficient peripheral myeloid cells occur in order to preserve lysosomal function and that this increase in GPNMB expression is associated with an anti-inflammatory environment and potentially dampened immune response.

## Methods

### Human Peripheral Blood Mononuclear Cells (PBMCs)

Samples from the National Centralized Repository for Alzheimer’s Disease and Related Dementias (NCRAD) were used. Cryopreserved peripheral blood mononuclear cells (PBMCs) from *GRN*-FTD and age- and sex-matched neurologically healthy controls (n=25 per participant group) were received and kept in liquid nitrogen cryostorage until use. See Supplementary File 1 for participant demographics.

### Human PBMC cryorecovery

For cryorecovery, PBMCs were retrieved from liquid nitrogen, thawed at 37°C, slowly added to 37°C filter sterilized complete culture media (RPMI 1640 media, 10% heat-inactivated fetal bovine serum (FBS), 1mM Penicillin-Streptomycin) and pelleted via centrifugation at 90 *x g* for 10 min at room temperature. The supernatant was removed and cells were resuspended in 37°C MACS buffer (PBS, 0.5% bovine serum albumin, 20 mM EDTA, pH 7.2) cell counting using Trypan blue exclusion.

### Nucleofection and plating of human PBMCs

Cells were aliquoted into 50 mL falcon tubes with 1 x 10^6^ cells per nucleofection reaction. Cells were centrifuged at 90 x *g* for 10 minutes at 4°C. The supernatant was carefully aspirated so as not to disturb the cell pellet, and cells were resuspended in nucleofection buffer (acclimated to room temperature; Lonza; P3 Primary Cell 4D-Nucleofector X Kit S; V4XP-3032) containing 1 µM *GPNMB* or control ASO per 20 µL (ASOs provided by Ionis, sequences detailed in Table 1), to a final concentration of 1 x 10^6^ cells per 20 µL. 20 µL of cells were transferred to each Nucleocuvette, which was then placed into a 4D-Nucleofector® X Unit (Lonza) and pulsed using the EO 115 pulse code. After nucleofection, 400 µL of growth media (acclimated in incubator 1-hour prior) was added to each Nucleocuvette and cells transferred to 24-well plates at a final concentration of 1 x 10^6^/mL, with 5 x 10^5^ cells per well. Cells were left to incubate for 24-hours, after which cells were stimulated with vehicle or 100U IFNγ (Peprotech) for 18-hours.

### Flow cytometry of human PBMCs

PBMCs were taken for flow cytometry and transferred to a v-bottom 96-well plate (Sigma, CLS3896-48EA) and centrifuged at 300 x *g* for 5 minutes at 4°C. Cells were resuspended in 50uL growth media containing Lysotracker™ Red DND-99 and BMV109 (Vergent Bioscience), both at 1:1000 dilution, and incubated at 37°C for 1-hour. Cells were centrifuged for 5 minutes at 300 x *g* at 4°C and washed in PBS x 2. Cells were then resuspended in 50 µL of PBS containing diluted fluorophore-conjugated antibodies (see Table 2) and incubated in the dark at 4°C for 20 minutes. Cells were centrifuged at 300*x g* for 5 minutes at 4°C and washed in PBS x 2. Cells were fixed in 50 µL of 1% paraformaldehyde (PFA) at 4°C in the dark for 30 minutes. Cells were centrifuged at 300*x g* for 5 minutes and resuspended in 200 µL FACs buffer (PBS, 0.5 mM EDTA, 0.1% sodium azide). Cells were taken for flow cytometry on a MACS Quant Analyzer (Miltenyi). A minimum of 100,000 events were captured per sample and data were analyzed using FlowJo version 10.6.2 software (BD Biosciences). When validating flow cytometry panels and antibodies, fluorescence minus one controls (FMOCs) were used to set gates and isotype controls were used to ensure antibody-specific binding.

### Cytokine release measurements via multiplexed immunoassays

V-PLEX mouse pro-inflammatory panel 1 kit (MesoScaleDiscovery; K15048D) or V-PLEX human pro-inflammatory panel 1 kit (MSD; K15049D) was used to quantify cytokines in conditioned media from pMacs. Media was diluted 1:1 with MSD kit diluent and incubated at room temperature in the provided MSD plate with capture antibodies for 2 hours as per manufacturer’s instructions. Plates were then washed x 3 with PBS with 0.05% Tween-20 and detection antibodies conjugated with electrochemiluminescent labels were added and incubated at room temperature for another 2 hours with shaking. After 3 x washes with PBS containing 0.05% Tween-20, MSD read buffer was diluted to 2x and added, and the plates were loaded into the QuickPlex MSD instrument for quantification. Results were normalized to total live cell counts as measured via flow cytometry.

### Animals

*Grn* knockout (*Grn* -/-) mouse strains have previously been characterized (*23*) and were maintained in the McKnight Brain Institute vivarium (University of Florida) at 22°C at 60-70% humidity and animals were kept in a 12-hour light/dark cycle. C57BL/6 littermate controls were used for all studies, with *Grn* -/- and C57BL/6 controls cohoused. All animal procedures were approved by the University of Florida Institutional Animal Care and Use Committee and were in accordance with the National Institute of Health Guide for the Care and Use of Laboratory Animals (NIH Publications No. 80-23) revised 1996. Male and female mice were aged to 5-6-months old and sacrificed via cervical dislocation or decapitation.

### Harvesting and culturing of peritoneal macrophages and ex-vivo stimulation of non-nucleofected cells

Peritoneal macrophages were harvested from mice which had received a 1-mL intraperitoneal administration of 3% Brewer thioglycolate broth 72-hours prior collection. Mice also received Buprenorphine Sustained-Release every 48-hours for pain relief. Mice were sacrificed via cervical dislocation and abdomen sprayed with 70% ethanol. The skin of the abdomen was split along the midline, taking care to avoid puncturing or cutting the abdominal cavity. 10 mL of cold RPMI media (Gibco; 11875119) was injected into the peritoneal cavity using a 27G needle. After gentle massaging of the peritoneal cavity, as much fluid was withdrawn as possible from the peritoneal cavity using a 25G needle and 10 mL syringe. Aspirated fluid was passed through a 70uM nylon filter onto 50mL falcon and pre-wet with 5mL of HBSS^-/-^. Filters were then washed twice with 5mL of HBSS^-/-^ and then tubes spun at 400 x *g* for 5 minutes at 4°C. Supernatant was aspirated and pellet resuspended in 3 mL pre-warmed growth media (RPMI, 10% FBS, 1% Pen-Strep). Cells were counted and viability recorded using trypan-blue exclusion on an automated cell-counter (Countess™; Thermo). Volume growth media was adjusted so that cells were plated at 5 x 10^5^/mL in 6-, 24- or 96-well plates depending on the intended assay. Cells were incubated at 37°C, 5% CO2 for a minimum of 3-hours to allow macrophages to adhere. Wells were washed twice with sterile PBS to remove non-adherent cells and new, pre-warmed growth media added. For cells requiring *ex-vivo* stimulation, 100U of IFNγ (R&D) or vehicle (H_2_O) was added for 18-hours. Protocol available at dx.doi.org/10.17504/protocols.io.j8nlkoyoxv5r/v1.

### Nucleofection and plating of mouse peritoneal macrophages

Peritoneal macrophages were harvested from mice as previously described. Once passed through 70uM nylon filter, tubes were centrifuged at 90 x *g* for 10 minutes at 4°C. Cells were resuspended and counted as previously described. Cells were aliquoted into 50 mL falcons with 1 x 10^6^ cells per nucleofection reaction. Cells were centrifuged at 90 x *g* for 10 minutes at 4°C. Supernatant was carefully aspirated so not to disturb the cell pellet, and cells resuspended in nucleofection buffer (acclimated to room temperature; Lonza; P2 Primary Cell 4D-Nucleofector X Kit L; V4XP-2024) containing 1 µM *GPNMB* or control ASO per 100 µL (sequences detailed in Table 1), to a final concentration of 1 x 10^6^ cells per 100 µL. 100 µL of cells were transferred to each Nucleocuvette, which was then placed into a 4D-Nucleofector® X Unit (Lonza) and pulsed using the CM 138 pulse code. After nucleofection, 400 µL of growth media (acclimated in incubator 1-hour prior) was added to each Nucleocuvette and cells transferred to plates pre-coated with poly-D-Lysine (Sigma). Cells were left to incubate for 24-hours, after which media was aspirated, cells washed and assays started as previously described.

### Flow cytometry of mouse peritoneal macrophages

1-hour prior to collection, BMV109 Pan Cathepsin probe (Vergent Bioscience) and DQ red BSA (Invitrogen) were added to each well at a final concentration of 1µM and 10µg/mL, respectively, and cells incubated at 37°C for 1-hour. Cells were then washed 3 times in sterile PBS, harvested, and transferred to a v-bottom 96-well plate (Sigma, CLS3896-48EA) and centrifuged at 300 x *g* for 5 minutes at 4°C. Cells were resuspended in 50 µL of PBS containing diluted fluorophore-conjugated antibodies (see Table 3) and incubated in the dark at 4°C for 20 minutes. Cells were centrifuged at 300 x *g* for 5 minutes at 4°C and washed in PBS x 2. Cells were fixed in 50 µL of 1% paraformaldehyde (PFA) at 4°C in the dark for 30 minutes. Cells were centrifuged at 300 x *g* for 5 minutes. For intracellular GPNMB staining, cells were resuspended in 100 µL of permeabilization buffer (eBiosciences, 00-8333-56) and incubated on ice for 15 min. Conjugated anti-GPNMB antibody (see Table 3) was added to each well and incubated on ice for 20 minutes. Cells were centrifuged at 300 x g for 5 minutes at 4°C and washed in PBS x 2 before being resuspended in 200 µL FACs buffer (PBS, 0.5 mM EDTA, 0.1% sodium azide). Cells were taken for flow cytometry on a Macs Quant Analyzer (Miltenyi) or BD LSR Fortessa™ Cell Analyzer. A minimum of 100,000 events were captured per sample and data were analyzed using FlowJo version 10.6.2 software (BD Biosciences). When validating flow cytometry panels and antibodies, fluorescence minus one controls (FMOCs) were used to set gates and isotype controls were used to ensure antibody-specific binding. experiments. Protocol is available at dx.doi.org/10.17504/protocols.io.rm7vzx9x4gx1/v1.

### DQ-BSA and BMV109 fluorescence microscopy

BMV109 Pan Cathepsin probe (Vergent Bioscience) and DQ red BSA (Invitrogen) were added to each well at a final concentration of 1µM and 10µG/mL, respectively, and cells were incubated at 37°C for 1-hour. Cells were washed 3 x DPBS^+/+^ and fixed for 10 minutes at room temperature in Invitrogen™ eBioscience™ Intracellular Fixation buffer. Cells were washed 3 x DPBS^+/+^ and incubated in 1 μg/ml DAPI (Life Technologies) for 10 minutes at room temperature in DPBS^+/+^. Cells were imaged using an EVOS™ M7000 (Invitrogen) at 20 x magnification. Image analysis was performed using Cellprofiler 4.2.5. Protocol available at dx.doi.org/10.17504/protocols.io.261gedk5ov47/v1.

### Ea(_52–68_) peptide loading assay

MHC II Ea chain (Ea) (52–68) peptide (Anaspec) was reconstituted in sterile distilled H_2_O to a final concentrate of 1mg/mL. Once peritoneal macrophages had adhered to plates, 5µg per well was added in growth media. Cells were incubated for 18-hours and taken forward for flow cytometry. Protocol available at dx.doi.org/10.17504/protocols.io.14egn3r3pl5d/v1.

### Immunoblotting

Media was aspirated and cells washed in PBS and lysed in RIPA buffer (50 mM Tris pH 8, 150 mM NaCl, 1% NP-40, 0.5% Na Deoxycholate, 0.1% SDS). Cell lysates were then centrifuged at 10,000 x *g* for 10 mins at 4°C. 6X Laemmli sample buffer added (Thermo Fisher) and samples were reduced and denatured at 95°C for 5 minutes. Samples were loaded into 4-20% Criterion Tris-HCl polyacrylamide gels (BioRad) alongside Precision Plus Protein Dual-Color Ladder (Biorad) to determine target protein molecular weight. Electrophoresis was performed at 100 V for ∼60 minutes and proteins transferred to a polyvinylidene difluoride (PVDF) membrane using a Trans-Blot Turbo Transfer System (BioRad) which utilizes Trans-Blot Turbo Midi PVDF transfer packs (BioRad) in accordance with manufacturer’s instructions. Prior to blocking, total protein was measured using Revert total protein stain (Licor) and imaged on the Odyssey FC imaging system (Licor). Membranes were then blocked in 5% non-fat milk in TBS/0.1% Tween-20 (TBS-T) for 1 hour at room temperate and subsequently incubated with primary antibody (see Table 4) in blocking solution overnight at 4°C. Membranes were washed with TBS-T (3 x 5 minutes) and incubated in horseradish peroxidase (HRP)-conjugated secondary antibody (1:1000) (BioRad) in blocking solution for 1 hour. Membranes were washed in TBS-T (3x 5 minutes) and developed using Super signal west femto/pico (Thermo). Membranes were imaged using the Odyssey FC imaging system and quantified using Image Studio Lite Version 5.2 (Licor).

### MHC-II antibody uptake assay

MHC-II uptake was adapted from (*39*). Briefly, cultured pMacs in 24-well plates were incubated in primary anti-MHC-II monoclonal antibody (mAB; BD Biosciences, see Table 5) at a final concentration of 5μg/mL for 20 minutes at 4°C in FACS buffer containing 10% normal goat serum, CD16/CD32 (Mouse BD Fc Block; BD Biosciences) at a final concentration of 1:100 and live/dead stain (see Table 5). Antibody solution was aspirated and cells were washed in ice-cold FACS buffer x 2. pMacs were incubated in prewarmed growth media at 37°C for 0, 1 or 2 hours to allow for the uptake of antibody-bound MHC-II complexes. “No endocytosis” controls were incubated in ice-cold FACS buffer and maintained on ice for 2 hours. At each time point, internalization was stopped by adding 1 mL of ice-cold FACS buffer. Antibody-bound MHC-II that was not internalized was labelled with a fluorescent secondary antibody (see Table 5) at a final concentration of 5μg/mL for 20 minutes on ice. Secondary antibody solution was aspirated and cells washed in ice-cold FACS buffer x 2. Cells were harvested and transferred to a v-bottom 96-well plate (Sigma, CLS3896-48EA) and centrifuged at 300 x *g* for 5 minutes at 4°C. Cells were fixed in 50 µL of 1% paraformaldehyde (PFA) at 4°C in the dark for 30 minutes. Cells were centrifuged at 300 x *g* for 5 minutes and resuspended in 200 µL FACs buffer. Cells were taken for flow cytometry on a Macs Quant Analyzer (Miltenyi) or BD LSR Fortessa™ Cell Analyzer. A minimum of 100,000 events were captured per sample and data were analyzed using FlowJo version 10.6.2 software (BD Biosciences).

### MHC-II recycling assay

MHC-II recycling was adapted from (*39*). Briefly, cultured pMacs in 24-well plates were incubated in primary anti-MHC-II monoclonal antibody (mAB; BD Biosciences, see Table 5) at a final concentration of 5μg/mL for 1 hour at 37°C in FACS buffer to allow for the antibody-binding and subsequent uptake of MHC-II complexes. Antibody solution was aspirated and 1mL of ice-cold FACS buffer added to each well to stop primary antibody uptake, and cells washed 2 x in ice-cold FACS buffer. To block any antibody-labelled MHC-II complexes that had not been internalized and was left on the plasma membrane, cells were incubated for 20 minutes on ice in FACS buffer containing goat anti-mouse IgG-Alexa 647 (see Table 5) at a final concentration of 5μg/mL. Cells were washed 2 x with ice-cold FACS buffer and complete growth media added to each well. Cells were returned to 37°C incubator for an additional 1, 3 or 18 hours. After each time point, 1 ml of ice-cold FACS buffer was immediately added to each well to stop recycling of MHC-II and cells washed x 2 with ice-cold FACS buffer. Cells were incubated for 20 minutes on ice in FACS buffer containing goat anti-mouse IgG-Alexa 488 (see Table 5) at a final concentration of 5μg/mL. This step will label any primary antibody-labelled MHC-II, that had been taken up in the first incubation step, evading labelling with goat anti-mouse IgG-Alexa 647 in the blocking step and had been recycled back to the plasma membrane. Cells were harvested and transferred to a v-bottom 96-well plate (Sigma, CLS3896-48EA) and centrifuged at 300 x *g* for 5 minutes at 4°C. Cells were fixed in 50 µL of 1% paraformaldehyde (PFA) at 4°C in the dark for 30 minutes. Cells were centrifuged at 300 x *g* for 5 minutes and resuspended in 200 µL FACs buffer. Cells were taken for flow cytometry on a Macs Quant Analyzer (Miltenyi) or BD LSR Fortessa™ Cell Analyzer. A minimum of 100,000 events were captured per sample and data were analyzed using FlowJo version 10.6.2 software (BD Biosciences).

### Quantification of GPNMB extracellular fragment via ELISA

Conditioned media was collected from the pMacs at the indicated times. After being transferred to a 1.5mL tube, the media was centrifugated at 10,000 x *g* for 10 minutes at 4°C to remove any cell debris. The supernatant was transferred to a new tube and stored at -20°C for later analysis. Endogenous GPNMB ECF was measured using a Biotechne GPNMB ELISA Kit (Biotechne/R&D Systems, DY2330) following manufacturer instructions with minor modifications. A four-parameter logistic (4-PL) curve was generated from the GPNMB ECF standards included in the kit. The standard curve was created with an online 4-PL curve calculator from AAT Bioquest (https://www.aatbio.com/tools/four-parameter-logistic-4pl-curve-regression-online-calculator). The equation from the 4-PL curve was used to determine the concentration of GPNMB ECF in the test samples.

### RNA extraction and qPCR of inflammatory genes and Gpnmb

RNA was isolated using a Qiagen Mini-kit (Qiagen, category number 74104), following manufacturer instructions with minimal alterations. The concentration of BME was increased to a final concentration of 20uM to inhibit endogenous RNase activity. After elution, RNA concentration was determined using a Denovix spectrophotometer. cDNA was generated using a High-Capacity cDNA Reverse Transcription Kit (Applied Biosystems™, catalog number 4374966) following kit instructions. 2x Universal SYBR Green Fast qPCR Mix (Abclonal, RK21203) was used with the indicated primers (see Table 6) for detection. A final mass of 6.25ng of cDNA was used in each well. Each sample was run in triplicate per gene. Variance above 0.4 standard deviations from the sample’s average were excluded.

### Statistics and data analysis

Data and statistical analyses were performed using IBM SPSS statistics 27 or GraphPad Prism 9. For assessing differences between groups, data were analyzed by either 1-way or 2-way analysis of variance (ANOVA), or by t-test. In instances when data did not fit parametric assumptions, Kruskal-Wallis non-parametric ANOVA was used. Post-hoc tests following ANOVAs were conducted using Tukey HSD or Bonferroni correction. Two-tailed levels of significance were used and p < 0.05 was considered statistically significant. Graphs are depicted by means +/- standard error of the mean (SEM).

## Results

### GPNMB ASO-mediated knockdown in FTD-GRN carrier monocytes increases lysosomal burden, HLA-DR expression and IL-1β secretion

Before assessing the effects of GPNMB knock-down on PBMC immune cell function and lysosomal homeostasis in FTD-*GRN* and control groups, the *GPNMB*-targeting ASO (Ionis) was first optimized in healthy control samples. PBMCs from neurologically healthy controls (NHCs) were nucleofected with 1, 2 or 5μg of control or *GPNMB-*targeting ASO per reaction (1x10^6^ cells per reaction) and assessed for GPNMB expression levels. A significant reduction in GPNMB median fluorescence intensity (MFI) was observed in all cell types nucleofected with 1μg *GPNMB*-targeting ASO relative to control ASO (Sup. Figure 1A-D). No significant effect of *GPNMB* ASO was observed at either of the higher concentrations, with these higher concentrations decreasing GPNMB even in control ASO conditions. To ensure that nucleofection did not have adverse effects on inflammatory responses or lysosomal function that may confound the interpretation of result, PBMCs from NHCs were nucleofected with 1μg control ASO per reaction (1x10^6^ cells per reaction) or left non-nucleofected to assess effects of nucleofection. Cells were then plated in the presence or absence of 100U IFNγ for 18 hours and inflammatory responses and lysosomal function assessed via flow cytometry. Nucleofection had no significant effect on live cell percentages, GPNMB expression, lysosomal function nor stimulation-dependent HLA-DR expression (Sup. Figure 1E-H). As no effects of nucleofection were observed, all experiments discussed here on directly compare the effects of *GPNMB* knock-down via ASO relative to control ASO, with no non-nucleofected control cells.

To assess the effects of *GRN* mutations on immune cell function and lysosomal homeostasis, and how loss of GPNMB may modulate these, PBMCs were collected from 25 FTD-*GRN* patients and 25 neurologically healthy, age- and sex-matched controls (NHCs) (See Sup. File 1 for participant demographics) and sourced from the National Centralized Repository for Alzheimer’s Disease and Related Dementias (NCRAD). PBMCs were thawed, nucleofected with 1μg of *GPNMB* or control ASO (1 x 10^6^ cells per reaction), plated and allowed to recover for 24 hours, after which, 100U IFNγ or vehicle control was added to each well and cells were incubated for 18h (Figure 1A). We first assessed GPNMB MFI in different PBMC immune cell subsets (Sup. Figure 2A) to assess effects of *GRN* mutations on GPNMB expression and efficacy of ASO-mediated *GPNMB* knock-down. It was observed that classical monocytes from FTD-*GRN* patients exhibited significantly increased GPNMB MFI in control ASO and vehicle conditions relative to NHCs (Figure 1B). Regarding other immune cell subtypes, GPNMB MFI was also significantly increased in B cells from FTD-*GRN* patients relative to NHCs (Sup. Figure 2B), however no significant effects of patient group were seen in T cells (Sup. Figure 2C). GPNMB knock-down decreased GPNMB MFI levels in both treatment conditions and patient groups in all cell types assessed.

**Figure 1.**
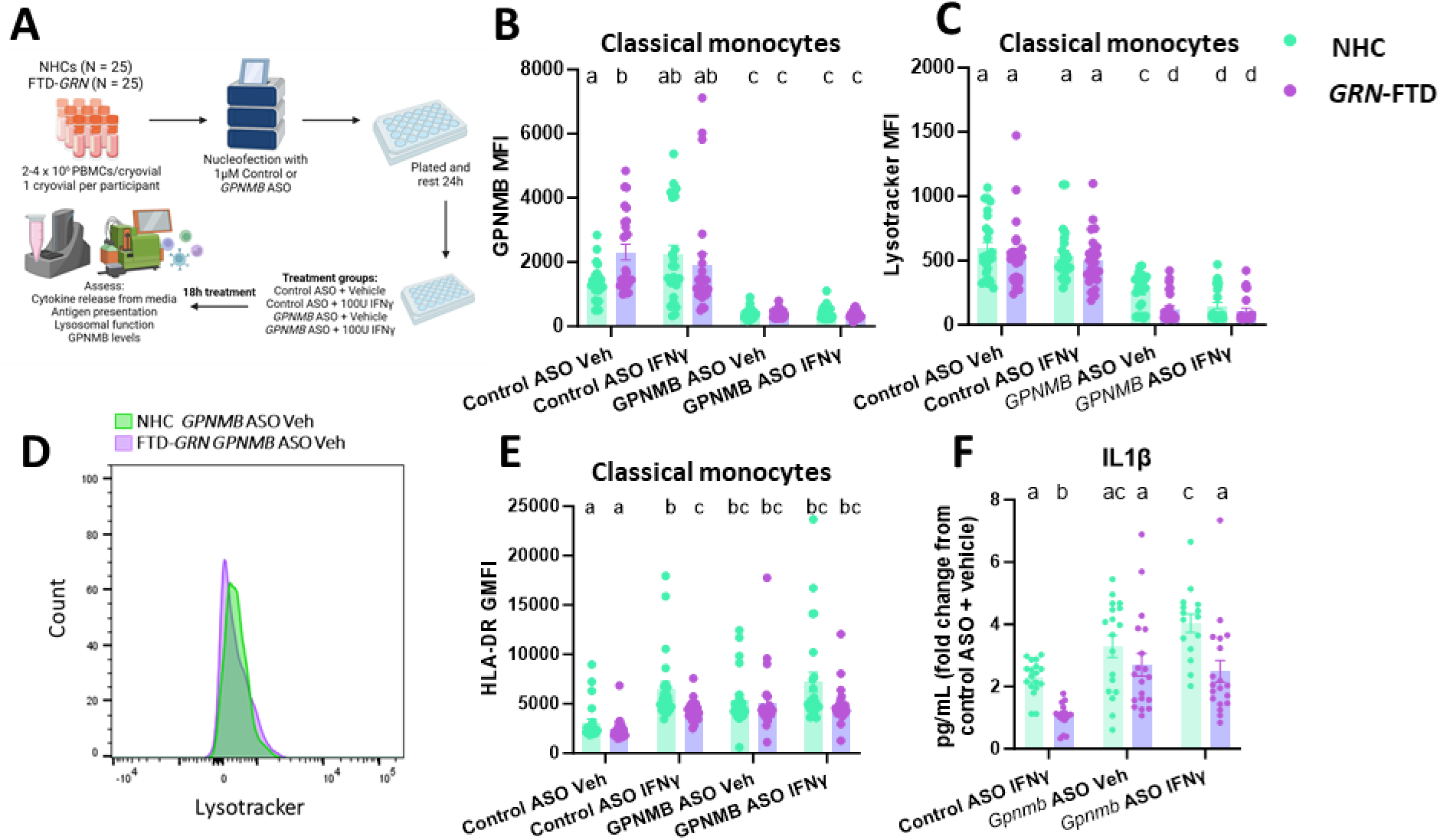
GPNMB ASO-mediated knockdown in FTD-*GRN* carrier monocytes increases lysosomal burden, HLA-DR expression and IL-1β secretion. PBMCs from NHCs and FTD-*GRN* patients were nucleofected with control or *GPNMB*-targeting ASO, plated and allowed to rest for 24 hours, followed by 18-hour incubation in presence or absence of 100U IFNγ and cells assessed via flow cytometry and media taken for cytokine quantification (**A**). GPNMB MFI was quantified in classical monocytes (**B**). Lysotracker MFI was quantified in classical monocytes (**C, D**). HLA-DR was quantified in classical monocytes (**E**). Cytokine release was quantified in media, normalized to live cell count and fold-change from control ASO vehicle conditions calculated (**F**). Bars represent mean +/- SEM (N = 20-25). Two-way ANOVA, Bonferroni post-hoc, groups sharing the same letters are not significantly different (p>0.05) whilst groups displaying different letters are significantly different (p<0.05).

Next, to assess the effects of *GRN* mutations and subsequent GPNMB knock-down on lysosomal function, MFI of Lysotracker DND-99, which will accumulate in and stain sufficiently acidic lysosomes within cells, was quantified in both classical monocytes and B cells. Surprisingly, no significant differences in Lysotracker MFI were exhibited between patient groups in control ASO conditions (Figure 1C). Interestingly, however, upon knock-down of GPNMB via ASO, a significant decrease in Lysotracker MFI was observed in FTD-*GRN* classical monocytes relative to NHCs in vehicle conditions (Figure 1C, D). This was not observed however in the presence of IFNγ. Surprisingly, no significant effects of participant group were observed in B cells in either control ASO or *GPNMB* ASO conditions (Sup. Figure 2D).

Given that many monocyte effector functions, such as antigen presentation and cytokine release, are dependent on lysosomal function, we next sought to assess the effects of *GRN* mutations and subsequent knock-down of GPNMB in these patient cells. When quantifying HLA-DR extracellular expression in classical monocytes, a significant decrease in stimulation-dependent HLA-DR expression was observed in FTD-GRN classical monocytes relative to NHCs in control ASO conditions (Figure 1E). Interestingly, with ASO-mediated knock-down of GPNMB, no stimulation-dependent increase in HLA-DR was observed in classical monocytes from either FTD-*GRN* patients or NHCs. This lack of response was due to a significant increase in HLA-DR expression in the absence of IFNγ in *GPNMB* ASO conditions relative to control ASO. It appears therefore, that an increase in GPNMB levels is associated with suppressed stimulation-dependent changes in HLA-DR expression, and that knock-down of GPNMB increases HLA-DR expression in classical monocytes. From this data alone, it is hard to determine if this increase in HLA-DR with GPNMB knock-down is driven by increased transcription and expression of HLA-DR, or if HLA-DR expression is increased at the plasma membrane due to altered internalization and recycling of the HLA-DR complex. Given the concomitant decrease in lysosomal function with the knock-down of GPNMB, the latter was hypothesized and was explored further in *ex vivo* murine cultures below.

Lastly, to determine if alterations in lysosomal function and HLA-DR expression were accompanied by alterations in cytokine release, we performed multiplexed immunoassays on conditioned media from plated PBMCs stimulated with vehicle or IFN-γ for 18 hours after nucleofection. When measuring cytokine secretion from PBMCs normalized to the percentage of live cells, taken from flow cytometry data previously described, no significant differences were detected between FTD-*GRN* patients and NHCs (Sup. Figure 2E-I). However, when these values were quantified as a stimulation-dependent fold-change from baseline (control ASO vehicle conditions), secretion of IL1β specifically was decreased in the FTD-*GRN* relative to NHCs in control ASO conditions (Figure 1F). Interestingly, upon knock-down of GPNMB, this fold-change was significantly increased in both patient groups, even in the absence of IFNγ, suggesting that GPNMB exerts anti-inflammatory effects, and knock-down of GPNMB increases pro-inflammatory milieu of PBMCs. Similar patterns were observed with TNF (Sup. Figure 2J), although no effect of GPNMB knock-down was observed. No significant effects of treatment or patient groups were observed for any other cytokine (Sup. Figure 2K-M).

### Grn-deficient mouse macrophages display upregulation of GPNMB which is responsive to inflammatory insults and ASO-mediated knockdown

To continue to investigate the effects of GPNMB knock-down in myeloid cells in the context of PGRN deficiency, assays were repeated in pMacs from *Grn* -/- mice and B6 controls. *GPNMB*-targeting and control ASOs were provided by Ionis. To optimize the use of these ASOs in pMacs, pMacs from 2-to-3-month-old B6 male mice were nucleofected with 2 different concentrations of *GPNMB* or control ASO (2 x 10^6^ cells per reaction) and GPNMB protein levels were assessed alongside non-nucleofected controls. A significant reduction in GPNMB protein levels were observed in pMacs nucleofected with both concentrations of *GPNMB*-targeting ASO, with no significant effect of control ASOs relative to non-nucleofected controls (Sup. Figure 3B, C). Moving forward, therefore, all experiments hereon were performed using the lower dose, at 1µg, of control or *GPNMB-*targeting ASO.

To ensure that nucleofection with this concentration of control ASO did not have adverse effects on inflammatory responses that may confound the interpretation of results, pMacs from 5-to-6-month-old, male and female, *Grn* -/- mice and B6 controls were nucleofected with control ASO and surface MHC-II expression at baseline, and in response to IFNγ, was quantified via flow cytometry in LPMs and SPMs, alongside non-nucleofected control cells. A significant main effect of treatment was observed, with IFNγ treatment significantly increasing surface MHC-II expression in both control ASO nucleofected and non-nucleofected LPMs and SPMs from both genotypes and sexes (Sup. Figure 3D-G). No significant differences were observed between non-nucleofected cells and those nucleofected with control ASO. Interestingly, a significant decrease in stimulation-dependent MHC-II expression was observed in LPMs from *Grn* -/- female mice relative to B6 controls, which was observed in both non-nucleofected and control ASO cells (Sup. Figure 3E). Furthermore, BMV109, a pan-cathepsin fluorescent probe, was used to quantify cathepsin activity in LPMs after nucleofection with a control ASO alongside non-nucleofected controls in order to assess effects of nucleofection on lysosomal function. No significant differences were observed between non-nucleofected and control ASO conditions in either genotypes or sexes (Sup. Figure 3H, I). Interestingly, a significant increase in BMV109 MFI was observed in LPMs from male *Grn* -/- mice relative to B6 controls which was unaffected by nucleofection (Sup. Figure 3H). Collectively, these data suggest that nucleofection with a control ASO at 1µg does not significantly modify immune responses or lysosomal function in macrophages relative to non-nucleofected cells. Therefore, all experiments discussed hereon directly compare the effects of GPNMB knock-down via ASO relative to control ASO, with no non-nucleofected control cells.

GPNMB expression was first assessed in pMacs from 5-to-6-month-old *Grn* -/- mice and B6 controls nucleofected with control or *GPNMB*-targeting ASO (Figure 2A). Both male and female mice were used and analyzed separately given the known sex-differences within the immune system (*40*). Nucleofected cells were plated and incubated with 100U IFNγ for 18-hours or a vehicle in order to assess effects of IFNγ on GPNMB expression. A significant increase in GPNMB expression was observed in both LPMs and SPMs from male and female *Grn* -/- mice relative to B6 controls in control ASO vehicle conditions (Figure 2B-E). Interestingly, IFNγ significantly reduced GPNMB expression in LPMs and SPMs from male and female *Grn* -/- from vehicle treated conditions to levels comparable to B6 controls. Such data suggests that GPNMB levels may be modulated by a pro-inflammatory environment, and GPNMB may increase to perturb inflammation in macrophages. ASO-mediated knock-down of GPNMB significantly reduced GPNMB expression in LPMs and SPMs in both sexes, genotypes and treatments. Orthogonal qPCRs were carried out to confirm our findings, and mRNA transcript levels of total pMacs reflected what was observed via flow cytometry, with the exception that, in control ASO pMacs treated with IFNγ, male *Grn* -/- mice retained an increase in *Gpnmb* expression relative to B6 controls, although it was observed to be significantly decreased from vehicle controls (Fig 2F, G). Collectively, this data demonstrates that GPNMB is elevated in LPMs and SPMs from 5-to-6- month-old, male and female, *Grn* -/- mice relative to B6 controls, and that GPNMB can be downregulated upon pro-inflammatory stimulus or ASO-mediated knock-down of GPNMB.

**Figure 2.**
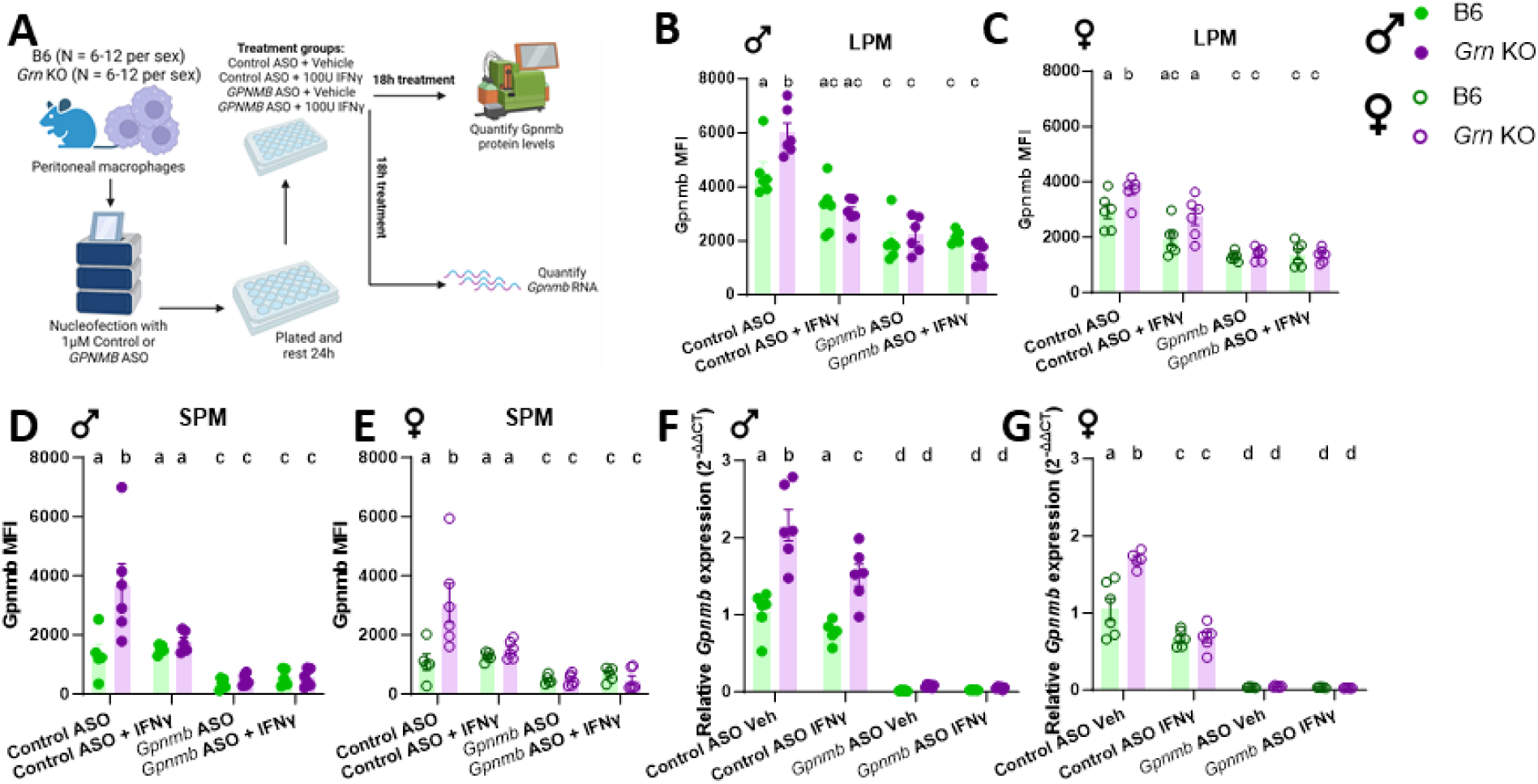
*Grn*-deficient mouse macrophages display upregulation of GPNMB which is responsive to inflammatory insults and ASO-mediated knockdown. pMacs from B6 and *Grn*-/-, male and female mice were nucleofected with control or *GPNMB*-targeting ASO, plated and allowed to rest for 24 hours, followed by 18-hour incubation in presence or absence of 100U IFNγ and cells assessed via flow cytometry or RNA extracted (**A**). GPNMB MFI was quantified in LPMs from male and female mice (**B, C**). GPNMB MFI was quantified in SPMs from male and female mice (**D, E**). Total GPNMB transcript levels were quantified in pMacs from male and female mice (**F, G**). Bars represent mean +/- SEM (N = 6-12). Two-way ANOVA, Bonferroni post-hoc, groups sharing the same letters are not significantly different (p>0.05) whilst groups displaying different letters are significantly different (p<0.05).

### ASO-mediated knockdown of GPNMB in Grn-deficient macrophages decreases lysosomal pan-cathepsin activity and protein degradation

In order to determine if, like in patient monocytes, GPNMB increases in *Grn* -/- macrophages to protect lysosomal function, nucleofected pMacs were incubated with the pan-cathepsin probe, BMV109, and DQ-BSA, a probe used to quantify lysosomal protein degradation, and fluorescence levels were quantified. Interestingly, in male pMacs nucleofected with control ASO, *Grn* -/- pMacs exhibited significantly increased BMV109 and DQ-BSA MFI, indicative of increased lysosomal function (Figure 3A-C). When GPNMB is knocked down via ASO, these lysosomal readouts are reduced to levels comparable to B6 pMacs, indicating that increased GPNMB levels in male *Grn* -/- pMacs contribute to increased lysosomal function. Interestingly, a different phenotypic pattern was observed in female *Grn* -/- pMacs. In female pMacs nucleofected with control ASO, no significant differences in BMV109 or DQ-BSA MFI were observed between genotypes (Figure 3D-F). However, much like what was observed in patient monocytes, a significant decrease in lysosomal protein degradation was observed in *Grn* -/- pMacs relative to B6 controls when GPNMB is knocked down via ASO. It seems, therefore, that in *Grn* -/- females, GPNMB increases to protect lysosomal function, and once knocked down, loss of GPNMB predisposes lysosomes to the dysfunction driven by PGRN deficiency. Importantly, manipulation of GPNMB expression via ASO did not affect lysosomal function in pMacs isolated from B6 mice of either sex, supporting our hypothesis that upregulated GPNMB expression supports lysosomal health in *Grn-*deficient pMacs in a compensatory mechanism.

**Figure 3.**
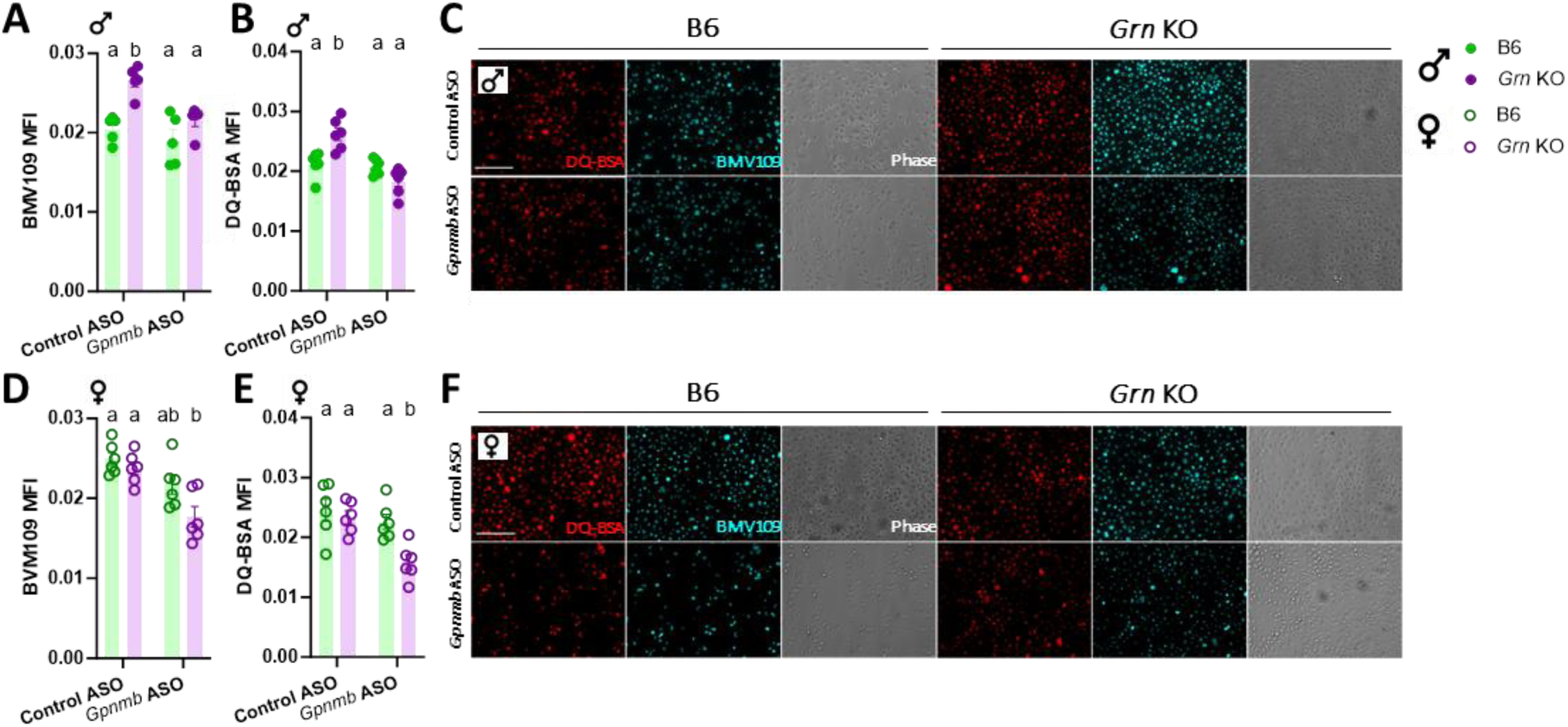
ASO-mediated knockdown of GPNMB in *Grn*-deficient macrophages decreases lysosomal pan-cathepsin activity and protein degradation. pMacs from B6 and *Grn* -/-, male and female mice were nucleofected with control or *GPNMB*-targeting ASO, plated and allowed to rest for 24 hours and lysosomal function assessed via microscopy. BMV109 and DQ-BSA MFI was quantified from microscopy images of pMacs from male mice (**A, B, C**). BMV109 and DQ-BSA MFI was quantified from microscopy images of pMacs from female mice (**D, E, F**). Bars represent mean +/- SEM (N = 6). Two-way ANOVA, Bonferroni post-hoc, groups sharing the same letters are not significantly different (p>0.05) whilst groups displaying different letters are significantly different (p<0.05). Scale bars, 30μM.

### ASO-mediated knockdown of GPNMB increases MHC-II surface expression, but not MHC-II processing in macrophages from Grn-deficient females

Next, to determine if alterations in lysosomal function were accompanied by alterations in MHC-II surface expression, nucleofected pMacs were treated with 100U IFNγ or vehicle for 18h, and surface MHC-II expression assessed on LPMs. Similar to what was observed in patient monocytes, in *Grn* -/- LPMs nucleofected with control ASO, a nonsignificant decrease in stimulation-dependent MHC-II expression was observed in males relative to B6 controls, and this decrease reached statistical significance in females (Figure 4A, B). Interestingly, when GPNMB is knocked down via ASO, an increase in stimulation-dependent MHC-II surface expression was observed in all genotypes and sexes. It seems, therefore, that with decreased levels of GPNMB, MHC-II expression increases. Indeed, correlational analysis demonstrated that GPNMB and MHC-II MFI negatively correlate, however this only reaches significance in *Grn* -/- pMacs (Figure 4C) suggesting an increased association between MHC-II and GPNMB in *Grn* deficient cells.

**Figure 4.**
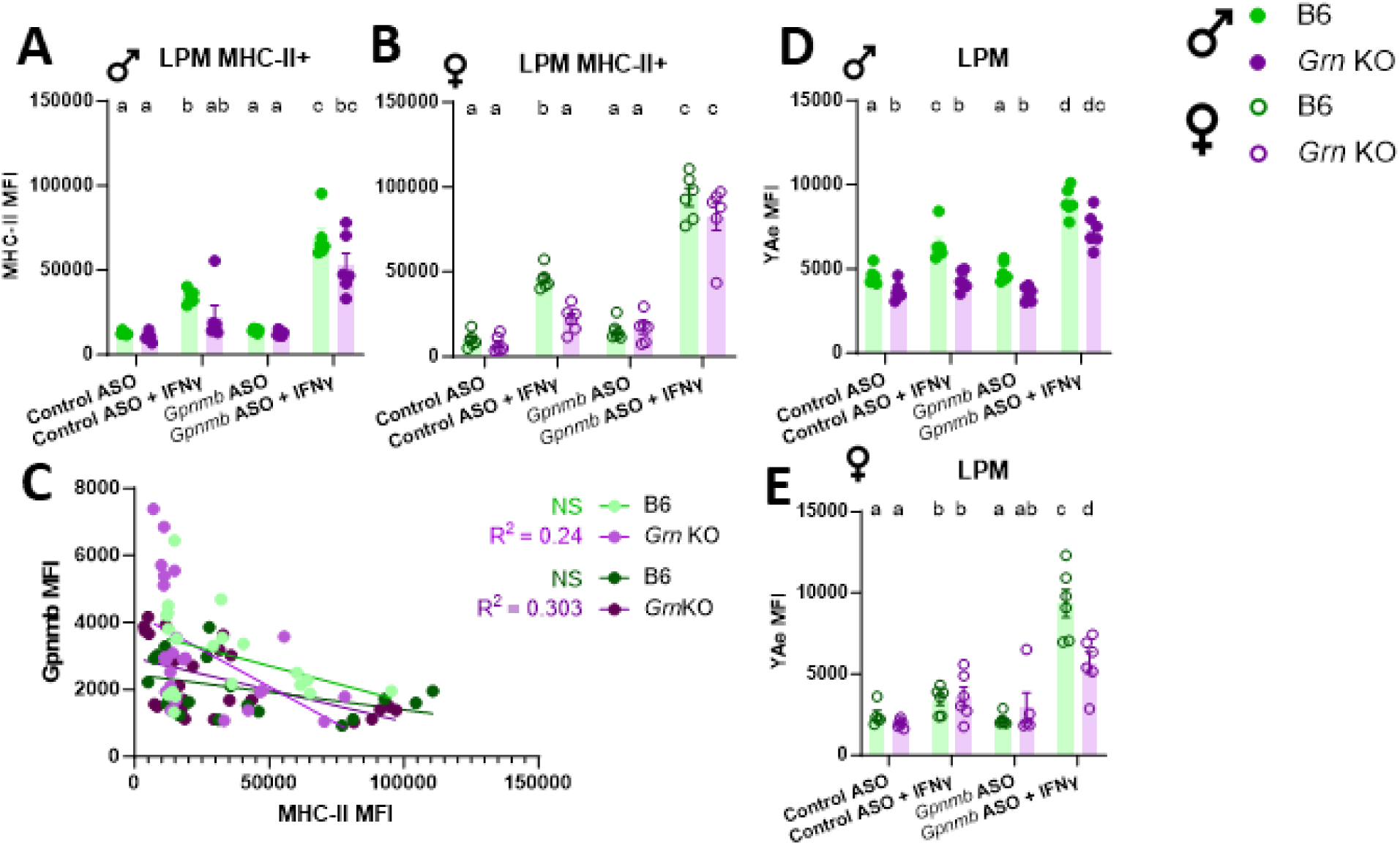
ASO-mediated knockdown of GPNMB increases MHC-II surface expression, but not MHC-II processing in macrophages from *Grn*-deficient females. pMacs from B6 and *Grn*-/-, male and female mice were nucleofected with control or *GPNMB*-targeting ASO, plated and allowed to rest for 24 hours, followed by 18-hour incubation in presence or absence of 100U IFNγ and cells assessed via flow cytometry. MHC-II MFI was quantified in MHC-II+ LPMs from male and female mice (**A, B**). Relationship between GPNMB and MHC-II MFI was assessed via correlational analysis in LPMs from male and female mice (**C**). Cells were incubated with Eα peptide and peptide-bound MHC-II complexes at the plasma membrane were quantified in LPMs from male and female mice (**D, E**). Bars represent mean +/- SEM (N = 6). Two-way ANOVA, Bonferroni post-hoc, groups sharing the same letters are not significantly different (p>0.05) whilst groups displaying different letters are significantly different (p<0.05).

What is perplexing is that antigen presentation and MHC-II complex assembly is highly dependent on lysosomal function. Yet, despite having decreased lysosomal function upon GPNMB knock down, female *Grn* -/- LPMs exhibit increased MHC-II surface expression. It may be, therefore, that MHC-II expression is increased on the surface of these cells due to altered processing and uptake of the MHC-II complex. The Eα: YAe model is used to monitor the antigen presentation capabilities of cells by incubating cells with an endogenous peptide (Eα 52–68) which is subsequently phagocytosed, transported to the lysosome, loaded onto an MHC-II complex at the lysosome and transported back to the plasma membrane for antigen presentation (Sup. Figure 4A). This Eα peptide-loaded MHC-II can subsequently be detected using flow cytometry using the YAe antibody (*38, 41*). This model allows us to measure antigen presentation of a peptide directly and acts as a measure of the whole antigen presentation pathway, from uptake to peptide loading to presentation. It was observed here that YAe MFI, in control ASO nucleofected LPMs from males, was significantly decreased in *Grn* -/- cells relative to B6 controls (Figure 4D). Upon GPNMB knock-down, however, stimulation-dependent Yae MFI significantly increased in both genotypes, with male *Grn* -/- LPMs exhibiting Yae MFI comparable to B6 control levels. Interestingly, no significant differences were observed between genotypes in control ASO nucleofected LPMs from females (Figure 4E). However, when GPNMB is knocked down, although significantly increasing from control ASO levels, LPMs from female *Grn* -/- mice exhibited decreased stimulation dependent YAe MFI relative to B6 controls, suggesting a sex-dependent decrease and dysregulation of the antigen processing pathway in these cells in the absence of GPNMB.

### ASO-mediated knockdown of GPNMB decreases MHC-II uptake and recycling in macrophages from Grn-deficient females GPNMB

Next, to further determine if increased MHC-II surface expression upon GPNMB knock down in female *Grn* -/- pMacs is due to altered MHC-II processing, uptake and recycling of MHC-II complexes were quantified via flow cytometry. MHC-II uptake can be quantified via flow cytometry by utilizing a previously described pulse-chase approach (*39*). Briefly, surface MHC-II complexes are labelled with a monoclonal primary antibody (pulse) and allowed to be internalized by the macrophage for 0, 1 or 2 hours (chase; Figure 5A). At each timepoint, surface MHC-II remaining can be detected via fluorescent secondary antibody. A significant decrease in surface MHC-II was seen over time as would be expected (Figure 5B). Control cells left on ice for 2 hours (no endocytosis control) showed no significant decrease in surface MHC-II, indicating no uptake of MHC-II. At both the 0- and 1-hour time point, no significant differences were observed between genotypes. However, at the 2-hour time point, a significant increase in MHC-II surface expression was observed when GPNMB is knocked down in *Grn* -/- pMacs from females relative to B6 controls and ASO control groups (Figure 5C). Such observations indicate that uptake of MHC-II complex from the plasma membrane is disrupted in the absence of GPNMB in *Grn* -/- female pMacs.

**Figure 5.**
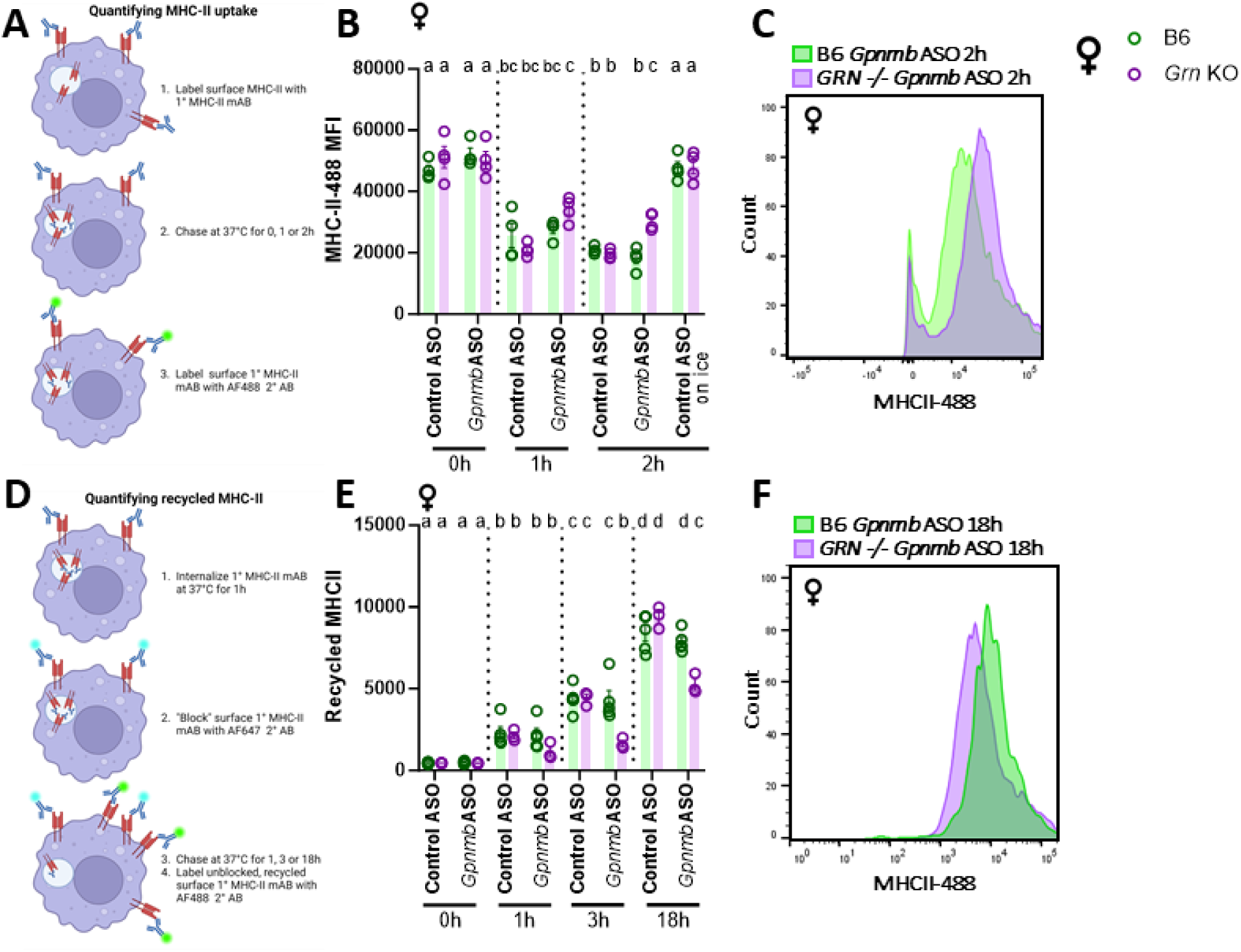
ASO-mediated knockdown of GPNMB decreases MHC-II uptake and recycling in macrophages from *Grn*-deficient females. pMacs from B6 and *Grn* -/- female mice were nucleofected with control or *GPNMB*-targeting ASO, plated and allowed to rest for 24 hours. After which, they were assessed for MHC-II uptake utilizing a pulse-chase flow cytometry-based assay (**A**). MHC-II-488 MFI was quantified in LPMs from female mice over a 2-hour time-course, with an ‘on ice’, no-endocytosis control included (**B, C**). pMacs were assessed for MHC-II recycling utilizing a pulse-chase flow cytometry-based assay (**D**). Recycled MHC-II MFI was quantified in LPMs from female mice over an 18-hour time-course (**E, F**). Bars represent mean +/- SEM (N = 6). Three-way ANOVA, Bonferroni post-hoc, groups sharing the same letters are not significantly different (p>0.05) whilst groups displaying different letters are significantly different (p<0.05).

Next, to determine if recycling of MHC-II back up to the cell surface once internalized is also impaired, we then used a pulse-chase approach that has previously been described (*42*). As with the uptake assay, surface MHC-II is first labelled with primary monoclonal antibody (Figure 5D) This labelled MHC-II is then allowed to be internalized for 1 hour at 37°C (pulse). Any remaining labelled surface MHC-II is then ‘blocked’ with secondary fluorescent antibody. Internalized, labelled MHC-II is then ‘chased’ for 1, 3 or 18 hours in the presence of IFNγ. At the end of each time-point, recycled MHC-II that has returned to the surface can be detected via fluorescent secondary antibody. A significant increase in recycled MHC-II expression is seen across timepoints as expected (Figure 5E). At the 0- and 1-hour timepoints, no significant differences are observed between genotypes or nucleofection conditions. However, at both the 3- and 18-hour time points, a significant reduction in recycled MHC-II is observed in female *Grn* -/- pMacs nucleofected with GPNMB-targeting ASO relative to B6 controls and control ASO conditions (Figure 5E, F). No significant differences between genotypes were seen in male pMacs, suggesting that, in males, the decrease in GPNMB does not significantly reduce MHC-II uptake and recycling in PGRN deficient cells as it does in females (Sup. Figure 5B, C).

### ASO-mediated knockdown of GPNMB dysregulates stimulation-evoked cytokine transcription and secretion normally mediated by the GPNMB ECF in mouse macrophages

Alterations in GPNMB expression have been shown to modulate cytokine expression and release in various immune cell subtypes (*24*). To determine how ASO-mediated knockdown of GPNMB would modulate cytokine levels in the context of PGRN deficiency, cytokine mRNA transcripts were quantified as well as cytokines secreted into cell culture media. In both female and male pMacs, a significant reduction in stimulation-dependent transcript levels of *Il6* were observed in *Grn* -/- pMacs in control ASO conditions (Figure 6A, B). Interestingly, upon knock-down of GPNMB, a significant increase in stimulation-dependent *Il6* production in both genotypes and sexes. However, despite this significant increase, *Il6* levels were still significantly decreased in *Grn* -/- pMacs nucleofected with GPNMB-targeting ASO relative to B6 controls. A similar phenotypic pattern was observed regarding IL6 secretion into the cell culture media (Figure 6C, D). Although a clear effect of genotype was not consistently observed in other cytokines quantified (Sup. Figure 6), a significant increase in cytokine transcript levels and/or production were observed upon the knock-down of GPNMB in both sexes and genotypes, suggesting that GPNMB may have an immunosuppressive effect on macrophage cytokine production and release.

**Figure 6.**
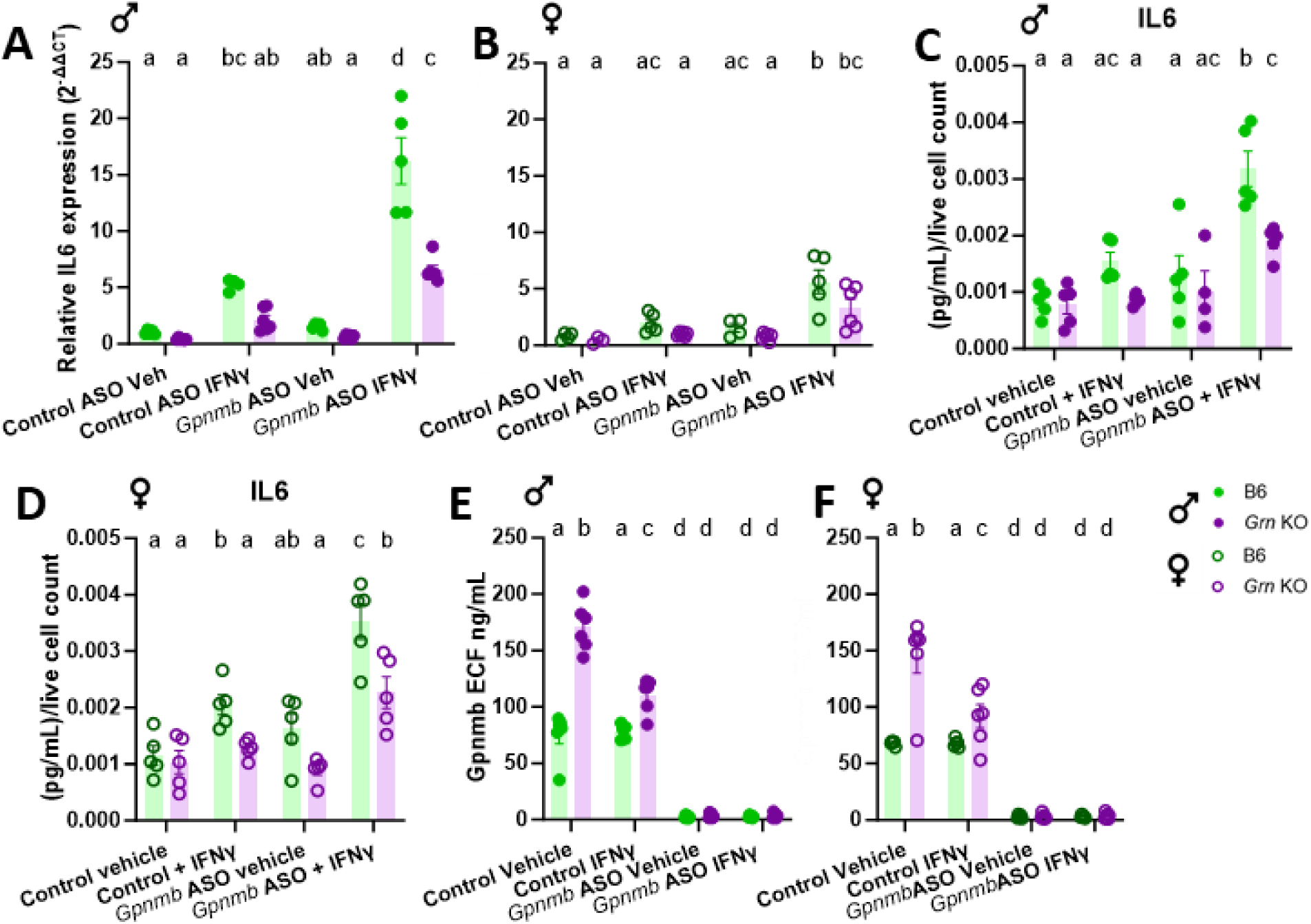
ASO-mediated knockdown of GPNMB dysregulates stimulation-evoked cytokine transcription and secretion normally mediated by the GPNMB ECF in mouse macrophages. pMacs from B6 and *Grn* -/- female and male mice were nucleofected with control or *GPNMB*- targeting ASO, plated and allowed to rest for 24 hours. After which, they were subject to 18-hour incubation in presence or absence of 100U IFNγ and cell RNA extracted and media collected. *IL6* transcript levels were assessed in pMacs from male and female mice (**A, B**). IL6 cytokine release in media was quantified in media and normalized to live cell count in pMacs from male and female mice (**C, D**). GPNMB ECF was quantified in media from pMacs from male and female mice (**E, F**). Bars represent mean +/- SEM (N = 5-6). Two-way ANOVA, Bonferroni post-hoc, groups sharing the same letters are not significantly different (p>0.05) whilst groups displaying different letters are significantly different (p<0.05).

GPNMB as a membrane protein can be cleaved and the extracellular fragment (ECF) acts as a paracrine factor to affect neighboring cells for modulating tumorigenesis and immunosuppression (*43*). We therefore hypothesized that the increase in cytokine production in pMacs nucleofected with *GPNMB*-targeting ASO could be driven by a decrease in the cleaved ECF being produced in the cell culture media. Indeed, when GPNMB ECF was quantified via ELISA in cell culture media, a significant reduction in *Grn* -/- pMacs was observed in both genotypes and sexes (Figure 6E, F). Interestingly, in control ASO conditions, GPNMB ECF levels reflected cellular protein expression that had been reported via flow cytometry (Figure 2), with significant increases observed in *Grn* -/- pMacs relative to B6 controls. Furthermore, GPNMB ECF is significantly reduced in control ASO nucleofected *Grn* -/- pMacs in the presence of IFNγ relative to vehicle controls, but still remained significantly elevated relative to B6 controls.

## Discussion

GPNMB expression is significantly upregulated in a number of different neurodegenerative diseases and our group has detected this upregulation in peripheral myeloid cells of young PGRN-deficient mice prior to any increases that have been reported in the CNS. However, why GPNMB increases in these diseases, and if it should be therapeutically targeted, is still up for debate. Here we demonstrate that, in both FTD-*GRN* patient monocytes and *Grn* -/- murine macrophages, GPNMB increases in the absence of PGRN, and GPNMB knockdown predisposes peripheral myeloid cells to lysosomal deficits. Furthermore, GPNMB upregulation is associated with an increase in internalization of surface MHC-II complexes and recycling, whereas a loss of GPNMB results in decreased uptake and less recycling, causing MHC-II complexes to accumulate on the cell surface. Also, increases in GPNMB cause a concomitant decrease in cytokine secretion and dampened response to pro-inflammatory stimulation, whereas GPNMB knock-down increases cytokine expression and secretion by myeloid cells.

GPNMB has been hypothesized to increase in the context of neurodegeneration due to lysosomal dysfunction. However, our findings herein suggest that GPNMB increases in both FTD-*GRN* carrier monocytes and *Grn* -/- mouse macrophages to protect lysosomal function, because the knock-down of it results in significant lysosomal dysfunction. It has previously been demonstrated that *Grn* -/- mice develop lipofuscinosis and lysosomal swelling in an age-dependent manner (*44–46*). Interestingly, lysosomal phenotypes have thus far only been reported in the brains of these mice at the earliest time-point of 10 months (*44*). It is therefore interesting and novel that we observed lysosomal deficits in *Grn* -/- macrophages from 5-to-6-month-old mice only upon GPNMB knock-down. As this is also the earliest time-point at which GPNMB upregulation has been reported in murine preclinical models (*25*), our data suggest that GPNMB upregulation is a compensatory mechanism to rescue and protect the lysosome in the absence of PGRN, and that this occurs much earlier in the peripheral immune system than in the CNS. As lysosomal deficits have been reported in aged animals, the upregulation of GPNMB to protect the lysosome may only be sufficient at these younger ages, and subsequent age-dependent phenotypes may render this protective mechanism insufficient.

Antigen presenting cells, such as macrophages, are highly dependent on lysosomal function for efficient peptide-loading and subsequent antigen presentation (*47–49*). The surface expression and transport of peptide-loaded MHC-II complexes (pMHC-II) are tightly regulated in macrophages and other antigen presenting cells. Specifically, newly synthesized MHC-II binds to a chaperone protein termed the Invariant chain (Ii) that remains associated with MHC-II during transport through the trans-Golgi network to the plasma membrane. Once at the plasma membrane, MHC- II–Ii complexes are rapidly internalized by clathrin-mediated endocytosis and traffic from early endosomes to late endosomal multivesicular antigen-processing compartments. It is in these compartments that MHC-II–associated Ii is degraded, immunogenic-peptides bind to MHC-II, and newly generated pMHC-II complexes traffic to the plasma membrane (*50*). Surface-expressed pMHC-II slowly internalizes, and like all internalized integral membrane proteins (*51*), internalized pMHC-II can either recycle back to the plasma membrane or traffic to lysosomes for degradation (*52*). Interestingly, we reported here for the first time that GPNMB upregulation in response to lysosomal dysfunction triggered by PGRN loss also mediates MHC-II complex cell surface expression and the ability of *Grn* -/- macrophages to engage in antigen presentation. With GPNMB knock-down, an increase in MHC-II extracellular expression was observed in *Grn* -/- macrophages, which was accompanied by a decrease in MHC-II uptake from the cell surface and a subsequent decrease in peptide loading and recycling to the plasma membrane. From our findings it is difficult to discern whether the accumulation of MHC-II at the cell surface upon GPNMB knock-down in *Grn* -/- macrophages represent MHC-II–Ii complexes or peptide-bound pMHC-II; however, given that a decrease in Eα peptide-bound MHC-II expression was observed in *Grn* -/- macrophages upon GPNMB knock-down, the former possibility is more likely, but further research is required to conclusively establish this. Alterations in MHC-II expression on macrophages would have deleterious consequences for subsequent macrophage effector function, specifically CD4+ and CD8+ T cell priming and induction of an effective immune response. It has been suggested that T cells are active players in neurodegeneration (*53*), consistent with reports of chemokine-driven recruitment of T cells observed in brain parenchyma of FTD patients relative to controls (*54*), and increased gene expression in genes associated with T cells in PBMCs from FTD subjects relative to controls (*55*). Future research is therefore required in order to understand the interplay between the innate and adaptive immune system in FTD-*GRN*, and how GPNMB might serve a protective role with respect to antigen presentation and macrophage effector function in early stages of disease pathogenesis.

Although it can be hypothesized that the upregulation of GPNMB in both FTD-*GRN* carriers and *Grn* -/- mouse macrophages is beneficial and occurs to ameliorate lysosomal deficits and allow sufficient MHC-II receptor uptake, recycling and processing through the lysosome, it is important to note that this was accompanied by reduced cytokine expression and release, suggesting an accompanying dampening or resolution of the immune response in these myeloid cells. A mechanism such as this one could prove to be beneficial, given previous reports that *GRN* mutations and deficiency cause increased pro-inflammatory cytokine release and expression (*23, 56*), which are thought to contribute to disease onset and/or progression (*57*). However, it is important to note that an excessively dampened immune response could be equally deleterious. For example, our group recently demonstrated that the PD-associated *R1441C-LRRK2* mutation induces immune cell exhaustion in both murine and human carrier monocytes (*37*). Interestingly, in light of the observation that brain-resident innate immune cells have been observed to increase communication with peripheral immune cells in response to an inflammatory insult, the CNS may heavily rely on peripheral immune cells to help curb peripheral inflammatory challenges that could adversely affect the brain (*58*). It has therefore been suggested that an exhausted peripheral immune system may be maladaptive, because it may be unable to respond to CNS signals to help curb inflammation in the CNS, thereby contributing to chronic neuroinflammation. Given that we observe a dampened immune response in association with increased GPNMB expression, we speculate that similar mechanisms may be contributing to neurodegeneration in the context of PGRN mutant/deficient patient cells, thereby representing an exciting avenue for future research. We demonstrated here that, although GPNMB protein and RNA levels are upregulated in pMacs from both male and female *Grn* -/- mice, knock-down of GPNMB only reveals lysosomal deficits and dysregulated MHC-II processing in pMacs from females. It seems, therefore, that male *Grn* -/- mice have additional compensatory mechanisms protecting lysosomal function which females do not possess. Interestingly, sex-specific differences have been reported regarding lysosomal function and gene expression (*59, 60*). Sex differences in the regulation of the autophagosome– lysosome system in particular have been proposed to modulate the severity of neurodegenerative disorders, because prior studies suggest that women have a lower basal autophagy (*61*). Such reports may explain why macrophages from female *Grn* -/- mice were vulnerable to lysosomal dysfunction when GPNMB was knocked down. Unlike Alzheimer’s Disease, which is overrepresented in females (*62*), FTD prevalence is found to be similar in men and women (*63*). However, sex has been found to influence clinical phenotype in FTD, with the behavioral variant of FTD reported to be more common in men, whereas primary progressive aphasia is overrepresented in women (*64*). The distribution of clinical phenotypes in the patient cohort assessed in this report were relatively evenly distributed between males and females (Table 7). Furthermore, this cohort was underpowered to stratify patient data by sex, therefore we are unable to comment on the effect of sex and clinical phenotypes on the cellular phenotypes reported here. However, whether differences in vulnerability of the lysosomal system is a mechanism underlying reported sex-differences in clinical phenotypes in FTD is of interest for future research.

GPNMB has been discussed as a potential therapeutic target in PGRN-mediated neurodegeneration (*21*). Based on the findings here, it appears that targeting GPNMB, specifically depleting GPNMB levels, would be deleterious as opposed to beneficial, given the protective effects of GPNMB on lysosomal function and antigen presentation observed here in *Grn* -/- macrophage and FTD-*GRN* patient monocytes. GPNMB has previously been suggested as a potential therapeutic target for interrupting the transmission of pathological α-synuclein in PD, due to the observation that GPNMB concentration in plasma positively correlates with UPDRS score, and knock-down of *GPNMB* in iPSC-derived neurons reduced α-synuclein pre-formed fibril (PFF) uptake (*65*). However, given the findings reported here, it could be argued that the positive correlation between GPNMB expression and UPDRS score may be driven by increased lysosomal dysfunction as disease progresses, with GPNMB increasing in an attempt to restore lysosomal function. Regarding altered PFF uptake, indeed, GPNMB knock-down may have reduced this, but this may have been due to decreased lysosomal function and altered endocytosis; such effects could arguably be equally deleterious in disease as PFF uptake. Given our observation that increased GPNMB expression was associated with reduced cytokine release, with knock-down ameliorating this, coupled with reports that GPNMB might have an anti-inflammatory role by promoting inflammation resolution (*24*), we propose a different strategy: that increasing GPNMB may be therapeutic for FTD and other neurodegenerative diseases where lysosomal function is impaired. However, as previously discussed, a dampened immune response may be equally as deleterious as an overactive one, and a suppressed immune response may even contribute to neurodegeneration (*37*).

### Conclusions

Our findings reported herein are consistent with a model in which GPNMB is upregulated in the absence of PGRN to protect lysosomal function, and with this comes a dampening of the immune system possibly to cap or resolve inflammatory challenges. Our results suggest that caution should be taken when targeting GPNMB for potential therapeutics, as reduction in GPNMB may put lysosomal function at risk and increase the pro-inflammatory milieu in the central-peripheral neuroimmune crosstalk which is critical for brain health (*66*). In fact, potential therapeutic strategies to increase GPNMB levels using small molecules or gene therapy may serve to rescue or protect the lysosome from further dysfunction and cap or curtail detrimental chronic inflammatory responses centrally and peripherally. In addition, the sex-dependent differences observed in preclinical animal models of disease suggest that future therapeutics should consider relative sex differences for more personalized medical approaches. Importantly, the associated curbing or dampening of immune responses triggered by increases in GPNMB should be investigated in more detail to determine the extent to which such a mechanism might contribute to neurodegeneration. Future research is needed to understand how the protective role that upregulation of GPNMB serves in the context of FTD-*GRN* mutant carriers interacts with an ageing immune system. Although we have demonstrated that upregulation of GPNMB in young, 5-to-6-month-old mice is associated with improved lysosomal function; how age modulates these phenotypes to eventually render them insufficient to prevent neurodegeneration is an unanswered question.

## List of abbreviations

PGRN: progranulin
FTD: frontotemporal dementia
LSD: lysosomal storage disorder
AD: Alzheimer’s Disease
PD: Parkinson’s Disease
CNS: central nervous system
GPNMB: glycoprotein non-metastatic B
ASO: antisense oligonucleotide
NCRAD: National Centralized Repository for Alzheimer’s Disease and Related Dementias
PBMCs: Peripheral blood mononuclear cells
NHCs: Neurologically healthy controls
pMHC-II: peptide-loaded MHC-II complexes

## Declarations

### Ethics approval and consent to participate

This study was reviewed and approved by the University of Florida Institutional Review Board (IRB202101359). All patient samples were obtained from NCRAD (see funding details below), and all participants provided informed consent in their respective cohort studies. This study was reviewed and approved by the University of Florida Institutional Animal Care and Use Committee (IACUC202200000114).

### Consent for publication

Not applicable

### Availability of data and materials

The datasets generated and analyzed during the current study are available in the Zenodo repository, 10.5281/zenodo.12773066

### Authors’ contributions

Conceptualization: RLW, MGT

Methodology: RLW, DAG, HAS, JM, SM, HK, NN

Funding acquisition: RLW, MGT

Project administration and supervision: MGT

Writing – original draft: RLW, HAS, DAG, JM, SM, NN, HK, MGT

All authors reviewed and approved the final manuscript.

## Acknowledgements

Regarding NCRAD samples, we thank contributors who collected samples used in this study, as well as patients and their families, whose help and participation made this work possible. We thank members of the Tansey lab for useful discussions and edits of the manuscript. We thank Biogen/Ionis for supplying all GPNMB and control ASOs. We thank the UF Interdisciplinary Center for Biotechnology Research (UF | ICBR) for use of flow cytometry facilities and advice.

