## Supplementary material for "ASO-mediated knockdown of GPNMB in mutant-*GRN* and *Grn*-deficient peripheral myeloid cells disrupts lysosomal function and immune responses": Sup materials and figures

### Supplementary figures

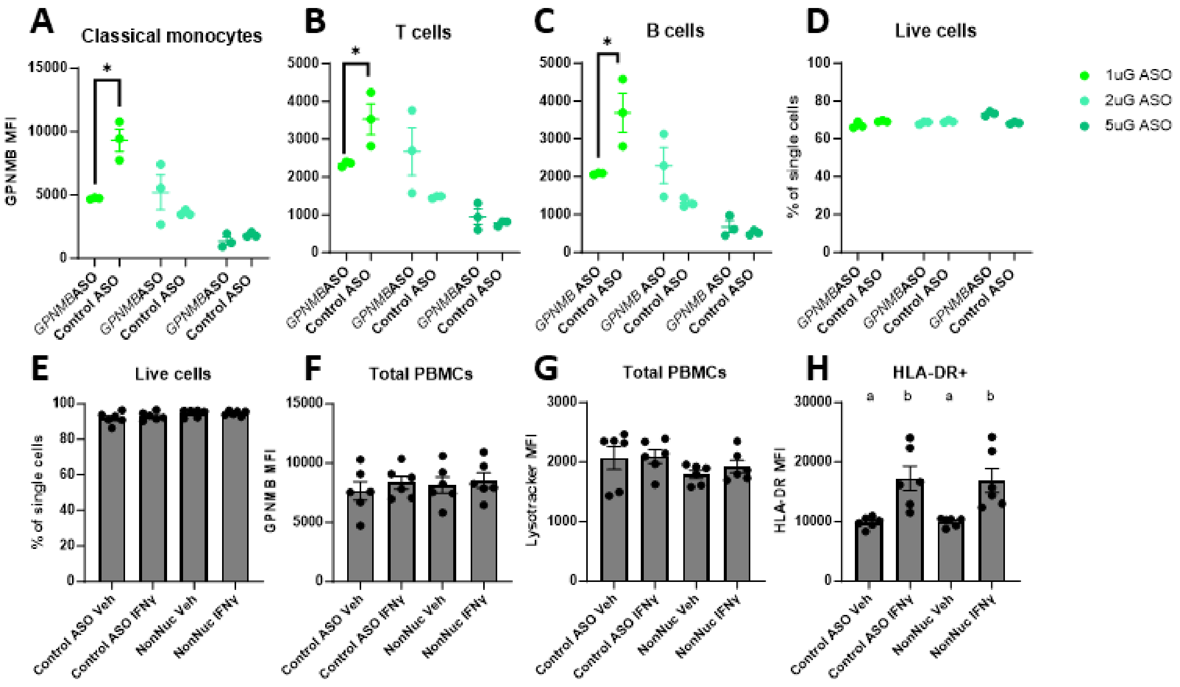


**Supplementary Figure 1. Optimization of GPNMB knock-down in human PBMCs.** PBMCs from NHCs were nucleofected with 1, 2 or 5µg *GPNMB*-targeting or control ASO and assessed for GPNMB MFI via flow cytometry (**A, B, C, D**). PBMCs from NHCs were nucleofected with 1µg control ASO or left unnucleofected and assessed for live cell frequency (**E**), GPNMB MFI (**F**), Lysotracker MFI (**G**) and HLA-DR MFI (**H**) via flow cytometry. Bars represent mean +/- SEM (N = 3-6) One/Two-way ANOVA, Bonferroni post-hoc, groups sharing the same letters are not significantly different ($p>0.05$) whilst groups displaying different letters are significantly different ($p<0.05$).

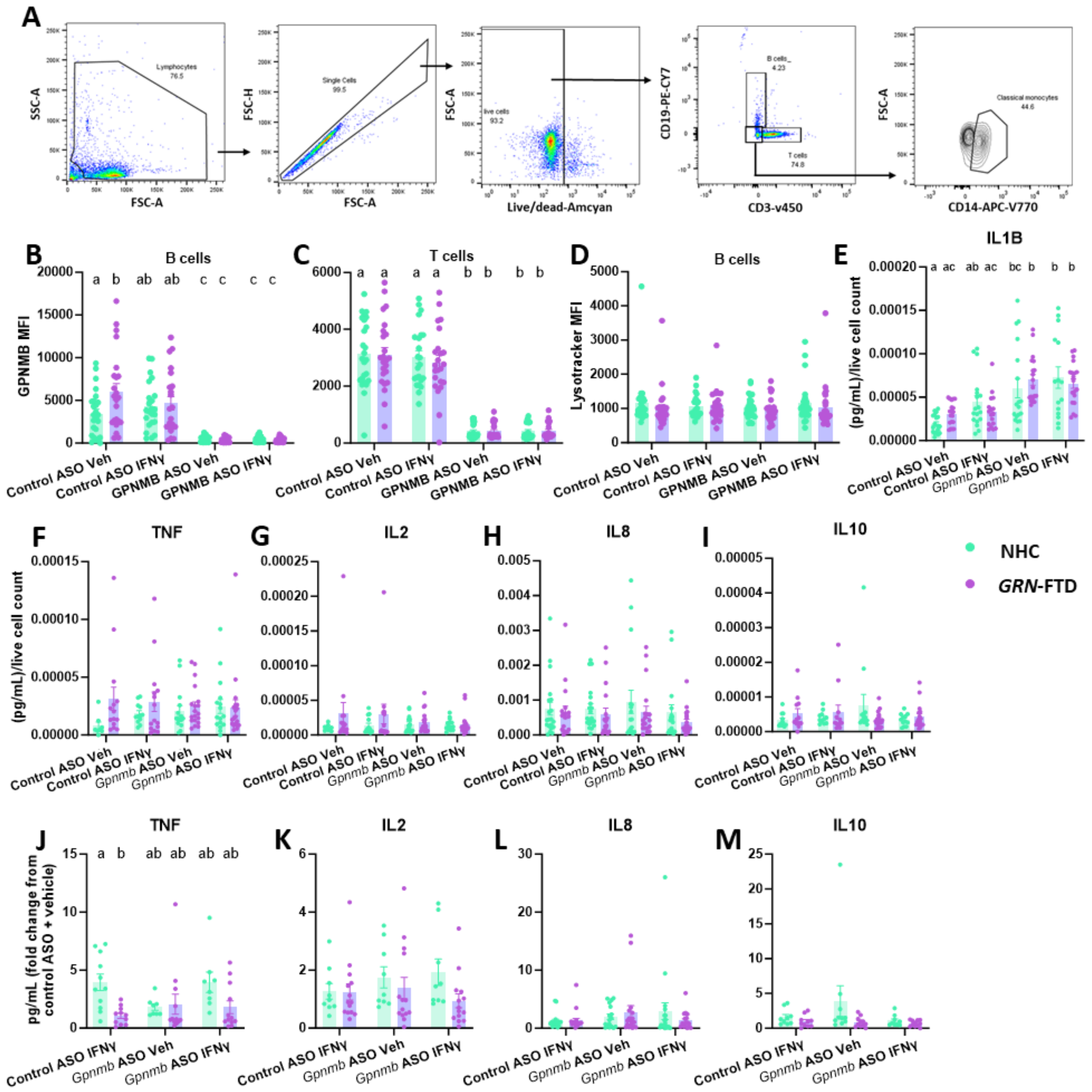


**Supplementary Figure 2. Immunophenotyping PBMCs and cytokine release.** PBMCs from NHCs and FTD-GRN patients were nucleofected with control or GPNMB-targeting ASO, plated and allowed to rest for 24 hours, followed by 18-hour incubation in presence or absence of 100U IFNγ and cells assessed via flow cytometry and media taken for cytokine quantification. (**A**) Schematic of flow cytometry gating strategy. GPNMB MFI was quantified in B and T cells (**B, C**). Lysotracker MFI was quantified in B cells (**D**). Cytokine release was quantified in media and normalized to live cell count (**E-I**). Cytokine release was quantified in media, normalized to live cell count and fold-change from control ASO vehicle conditions calculated (**J-M**). Bars represent mean +/- SEM (N = 20-25) Two-way ANOVA, Bonferroni post-hoc, groups sharing the same letters are not significantly different ($p>0.05$) whilst groups displaying different letters are significantly different ($p<0.05$).

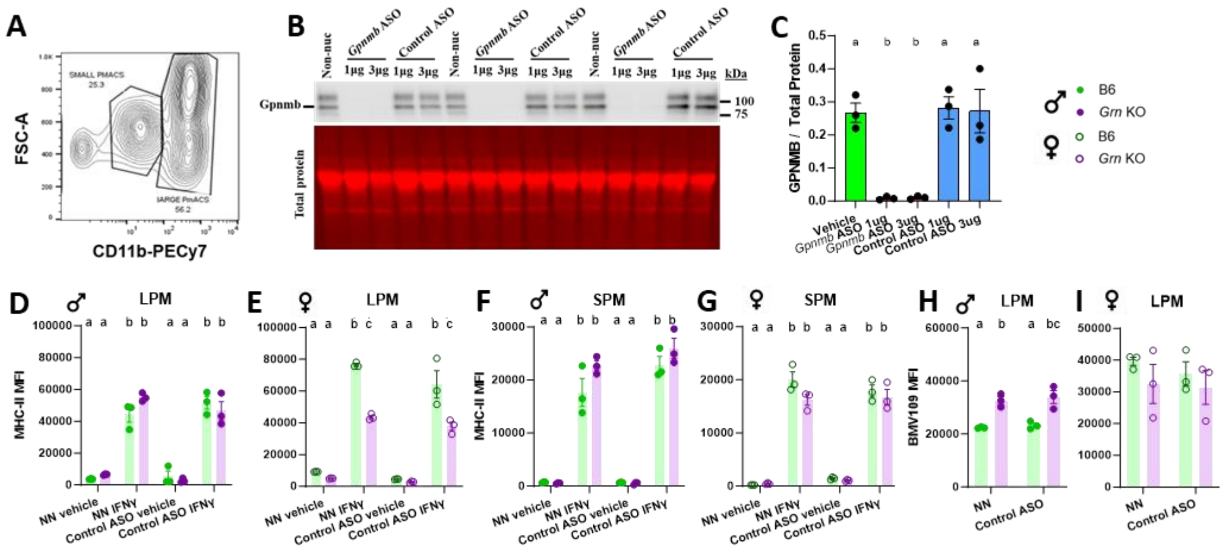


**Supplementary Figure 3. Optimization of GPNMB knock-down in murine pMacs.** (**A**) Schematic of pMac gating strategy to distinguish LPM and SPM based on Cd11b expression. pMacs from B6 mice were nucleofected with 1 or 3µg control or *Gpnmb*-targeting ASO or left unnucleofected and Gpnmb protein expression assessed via western blot (**B, C**). pMacs from B6 and *Grn* -/- mice were nucleofected with 1µg control ASO or left unnucleofected and assessed for MHC-II MFI and lysosomal function via flow cytometry (**D-I**). Bars represent mean +/- SEM (N = 3) One/Two-way ANOVA, Bonferroni post-hoc, groups sharing the same letters are not significantly different ($p>0.05$) whilst groups displaying different letters are significantly different ($p<0.05$).

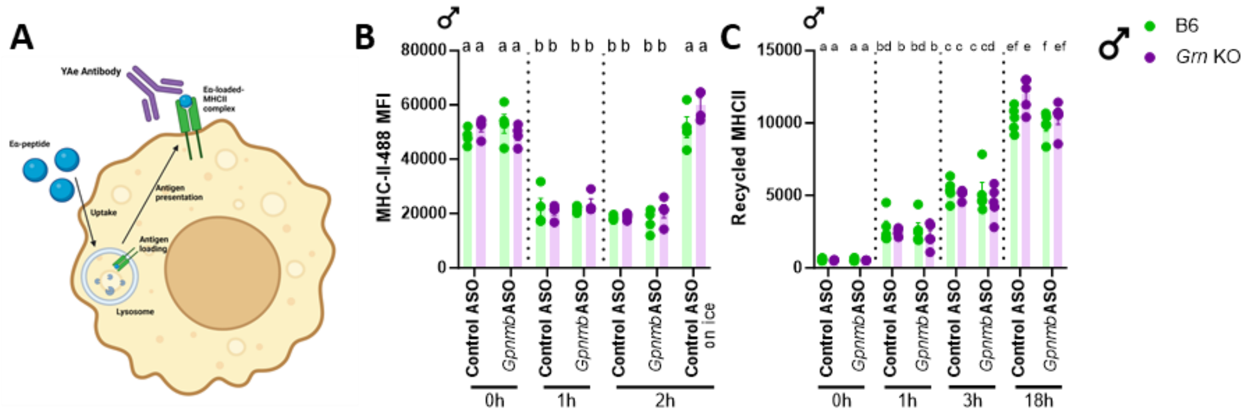


**Supplementary Figure 4. MHC-II processing in pMacs.** (**A**) Schematic of YAe flow-cytometry based assay**.** pMacs from B6 and *Grn* -/- male mice were nucleofected with control or *Gpnmb*-targeting ASO, plated and allowed to rest for 24 hours. After which, they were assessed for MHC-II uptake utilizing a pulse-chase flow cytometry-based assay. MHC-II-488 MFI was quantified in LPMs from male mice over a 2-hour time-course, with an 'on ice', no-endocytosis control included (**B**). pMacs were assessed for MHC-II recycling utilizing a pulse-chase flow cytometry-based assay. Recycled MHC-II MFI was quantified in LPMs from male mice over an 18-hour time-course (**C**). Bars represent mean +/- SEM (N = 6). Three-way ANOVA, Bonferroni post-hoc, groups sharing the same letters are not significantly different (p>0.05) whilst groups displaying different letters are significantly different (p<0.05).

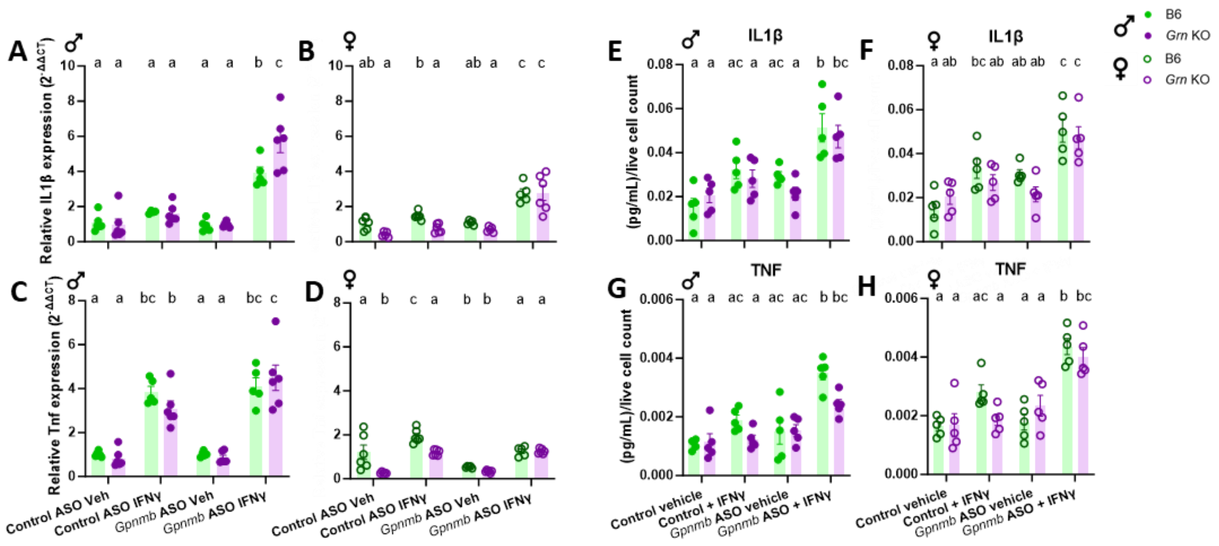


**Supplementary Figure 5. GPNMB regulates cytokine transcription and secretion in macrophages.** pMacs from B6 and *Grn* -/- female and male mice were nucleofected with control or *Gpnmb*-targeting ASO, plated and allowed to rest for 24 hours. After which, they were subject to 18-hour incubation in presence or absence of 100U IFNγ and cell RNA extracted and media collected. *IL1β and Tnf* trasncript levels were assessed in pMacs from male and female mice (**A, B, C, D**). IL1β and Tnf cytokine release in media was quantified in media and normalized to live cell count in pMacs from male and female mice (**E, F, G, H**). Bars represent mean +/- SEM (N = 5-6). Two-way ANOVA, Bonferroni post-hoc, groups sharing the same letters are not significantly different ($p>0.05$) whilst groups displaying different letters are significantly different ($p<0.05$).

### Tables

| ASO ID | Target | Sequence |
|---|---|---|
| 1391805-4 | mouse GPNMB | CTGTTTCCAAATTTCCCACA |
| 1392176-3 | human GPNMB | GCTACATTCTTTCTTGGACC |
| 676630 | Control ASO | CCTATAGGACTCTCCAGGAA |

**Table 1. ASO sequences**

| Target | Conjugate | Antibody Cat# | Dilution | Company |
|---|---|---|---|---|
| CD3 | V450 | 560365 | 20 | BD biosciences |
| Live dead | Amcyan | L34957 | 2000 | Invitrogen |
| HLA-DR | BV650 | 564231 | 20 | BD biosciences |
| GPNMB | PE | 12983842 | 20 | eBioscience |
| CD19 | PE Cy7 | 560728 | 50 | BD biosciences |
| CD14 | APCv770 | 130-098-076 | 20 | Miltenyi |
| FcX block | | 422302 | 20 | Biolegend |

**Table 2. Human PBMC flow cytometry antibody panel**

| Target | Conjugate | Antibody Cat# | Dilution | Company |
|---|---|---|---|---|
| Live dead | Amcyan | L34957 | 2000 | Invitrogen |
| FcR block | | 14-0161-85 | 100 | eBioscience |
| Gpnmb | CoraLite® 594 | CL59466926 | 100 | ThermoFisher |
| Cd11b | PE-Cy7 | 101216 | 200 | BioLegend |
| MHCII | APC-Cy7 | 107628 | 100 | BioLegend |
| Yae | FITC | 11-5741-82 | 100 | ThermoFisher |

**Table 3. Mouse pMac flow cytometry antibody panel**

| Target | Antibody Cat# | Dilution | Company |
|---|---|---|---|
| GPNMB | Ab188222 | 1000 | Abcam |
| Revert Total Protein | 926-11011 | - | Licor |

**Table 4. Western blotting antibodies**

| Assay | Primary or secondary | Target | Conjugate | Antibody Cat# | Dilution | Company |
|---|---|---|---|---|---|---|
| MHC-II uptake | Primary | MHC-II | - | MCA46GA | 100 | BioRad |
| | Secondary | - | AF488 | A11001 | 200 | Invitrogen |
| MHC-II recycling | Primary | MHC-II | - | MCA46GA | 100 | BioRad |
| | Secondary (block residual surface MHCII) | - | AF647 | A21237 | 200 | Invitrogen |
| | Secondary (label recycled MHCII) | - | AF488 | A11001 | 200 | Invitrogen |

**Table 5. Antibodies for MHCII uptake and recycling assays**

| Primer | Sequence |
|---|---|
| IL1B forward | CAA CCA ACA AGT GAT ATT CTC CAT G |
| IL1B reverse | GAT CCA CAC TCT CCA GCT GCA |
| IL6 forward | CAC AAG TCG GAG GCT TAA T |
| IL6 reverse | AAT TGC CAT TGC ACA ACT C |
| TNF forward | CTG AGG TCA ATC TGC CCA AGT AC |
| TNF reverse | CTT CAC AGA GCA ATG ACT CCA AAG |
| GPNMB forward | AAG CGA TTT CDG GATGTG CT |
| GPNMB reverse | CTT CCC AGG AGT CCTTCC AC |

**Table 6. Primer sequences for RTqPCR**

| | Female frequency | Male frequency |
|---|---|---|
| **bvFTD** | 57.1 | 45.5 |
| **PPA** | 14.3 | 9.1 |
| **CBS** | 14.3 | 27.3 |
| **MCI** | 7.1 | 9.1 |
| **AD** | 7.1 | 9.1 |

**Table 7. Distribution of clinical phenotypes between male and female patients.** bvFTD = Behavioral variant frontotemporal dementia, PPA = Primary progressive aphasia, CBS = Corticobasal syndrome, MCI = Mild Cognitive Impairment, AD = Alzheimer's Disease.
